# Parkinson’s Disease-Associated, Sex-specific Changes in DNA Methylation at *PARK7* (DJ-1), *ATXN1, SLC17A6, NR4A2*, and *PTPRN2* in Cortical Neurons

**DOI:** 10.1101/2021.09.08.459434

**Authors:** Joseph Kochmanski, Nathan C. Kuhn, Alison I. Bernstein

**Affiliations:** Department of Translational Neuroscience, College of Human Medicine, Michigan State University, Grand Rapids, MI, United States

**Author notes:** **Corresponding author:** Alison I. Bernstein, Department of Translational Neuroscience, College of Human Medicine, Michigan State University, Grand Rapids, MI, United States.

## Abstract

Evidence for epigenetic regulation playing a role in Parkinson’s disease (PD) is growing, particularly for DNA modifications. Approximately 90% of PD cases are due to a complex interaction between age, genes, and environmental factors, and epigenetic marks are thought to mediate the relationship between aging, genetics, the environment, and disease risk. To date, there are a small number of published genome-wide studies of DNA modifications in PD, but none accounted for cell-type or sex in their analyses. Given the hetereogeneity of bulk brain tissue samples and known sex differences in PD risk, progression, and severity, these are critical variables to account for. In this first genome-wide analysis of DNA methylation in an enriched neuronal population from PD post-mortem parietal cortex, we report sex-specific PD-associated methylation changes in *PARK7* (DJ-1), *SLC17A6* (VGLUT2), *PTPRN2* (IA-2β), *NR4A2* (NURR1), and other genes involved in developmental pathways, neurotransmitter packaging and release, and axon and neuron projection guidance.

## Introduction

Parkinson’s disease (PD) is the second most common neurodegenerative disorder in the US, and is characterized by progressive degeneration of dopaminergic neurons in the nigrostriatal pathway and the formation of α-synuclein-containing Lewy bodies.^1^ An estimated 5-10% of PD cases are familial and several genes have been identified that cause these inherited forms of the disease.^2,3^ The remaining ~90% of sporadic cases are likely due to a complex interaction between age, genes, and environmental factors. Given that epigenetic marks are dynamic with age, sensitive to the environment, and regulate gene expression throughout the lifespan, they are considered a potential mediator of the complex relationship between aging, genetics, the environment, and disease.^4,5^

Evidence for a role of epigenetic regulation in PD is growing, particularly for DNA modifications.^6–12^ In particular, previous studies have reported differential DNA methylation at PD-related genes (*MAPT, CYP2E1*, and *STX1B*),^13–15^ and hypomethylation of the α-syn gene (SNCA) is associated with decreased protein levels in the substantia nigra (SN) and striatum of post-mortem PD brain.^16–23^ α-syn also sequesters DNA methyltranserase 1 in the cytosol, away from the nucleus, resulting in global hypomethylation in cells overexpressing α-syn.^18^ In addition to these gene-specific studies, genome-wide analyses of DNA methylation from post-mortem PD brain tissue have identified a number of gene regions that show differential DNA methylation in PD brains.^24–27^ Finally, a case-control study identified an association between a polymorphism in PD risk and DNA methyltransferase 3B (DNMT3B), which is responsible for establishing de novo patterns of DNA methylation during embryonic development.^28^ Taken together, these multiple lines of evidence support a role for dysregulation of DNA methylation in the etiology of PD.

However, these previous genome-wide analyses of DNA modifications in PD have not adequately addressed the effects of sex or cell-type heterogeneity. Given that men and women show differences in PD risk, disease progression, and disease severity,^29–31^ it is critical to include sex in experiments examining the potential role for the epigenome in PD. Furthermore, the majority of existing studies of epigenetics in PD have utilized either blood or bulk brain tissue samples.^24,26,32,33^ Although DNA methylation changes in blood are useful for identifying biomarkers, they may not be representative of a pathogenic mechanism in the brain. Further, analyses from bulk brain tissue cannot address the cell-type specificity of identified changes, making it difficult to determine which cell types may be driving PD-related differential DNA methylation. In a step forward for the field, one recent paper investigated DNA methylation in an enriched neuronal population using fluorescence-activated cell sorting, but only explored DNA methylation at enhancer regions, not genome-wide.^27^ Here, we report the first genome-wide analysis of DNA methylation in enriched neurons from PD brain stratified by sex.

In this study, we obtained human postmortem parietal cortex from the Banner Sun Health Research Institute Brain Bank and enriched for neuronal populations using magnetic-activated cell sorting (MACS) for the neuronal marker, NeuN. Genome-wide DNA modifications were measured using the Illumina EPIC BeadChip array paired with bisulfite treatment. Bisulfite treatment is a well-established method for measuring DNA modifications, but it cannot distinguish between DNA methylation and DNA hydroxymethylation.^34,35^ As such, our dataset actually measures both DNA methylation and DNA hydroxymethylation without discriminating between these two epigenetic marks. Despite this important caveat, for simplicity’s sake and to match prior literature, we will discuss our results as changes in “DNA methylation.”

In our analysis, we identified sex-specific alterations in DNA methylation associated with PD. The most significant site for males is located within the *PARK7* (DJ-1) locus and the most significant site for females within the *SLC17A6* gene (VGLUT2). We also identified a region at the *PTPRN2* (Protein Tyrosine Phosphatase Receptor Type N2; IA-2β) locus that showed altered DNA methylation in both sexes, though the direction of change was sex-specific. In both the individual cytosine and regional analyses, we found almost no overlap of differential methylation between sexes, suggesting that PD-related changes in DNA methylation are sex-specific.

We also tested the hypothesis that PD is associated with accelerated epigenetic aging in the brain. Aging is the primary risk factor for PD, and previous work has shown that aging and PD progress via shared cellular mechanisms.^36–39^ Multiple hypotheses have been proposed to explain the association between aging and development of PD, including the ‘multiple hit’ hypothesis and the ‘stochastic acceleration hypothesis.’^37,40^ In general, these hypotheses work under a similar biological model – that an accumulation of factors, both genetic and environmental, accelerate the normal pace of dopaminergic neuron loss with age, eventually exceeding a threshold for PD diagnosis.^37^ However, the biological mechanism by which environment and genetics interact to accelerate age-related dopaminergic neuron loss remains unclear. Because the epigenome is dynamic with age and shows programmed age-related changes, it has been proposed that acceleration of these age-related changes can contribute to disease risk in the aged human.^41–44^ Supporting this idea, studies have shown associations between accelerated epigenetic aging and a variety of disease states, including a study using blood samples from PD patients.^32,45^ However, to our knowledge, no studies have investigated whether neurons show altered epigenetic aging in PD brains compared to control. Our epigenetic clock analysis showed no accelerated epigenetic aging in cortical neurons from PD brains compared to control.

## Results

### Human brain tissue sample selection

Control (n=50) and Parkinson’s disease (n=50) human brain specimens were obtained from archival human autopsy specimens provided by the Banner Sun Health Research Institute (BSHRI), using approved IRB protocols. Further details about the BSHRI’s brain samples are available in a previous publication.^46^ We selected PD patients with mid-to-late stage disease (Braak stage ≤ IV), as defined by Lewy pathology.^47^ The cohort of control brains consisted of patients who died from non-neurologic causes and whose brains had no significant neurodegenerative disease pathology. Subjects were split by sex (n=63 males, n=37 females), and data for the sample cohort is summarized in Table 1. For each subject (N=100), parietal cortex was obtained, as this region develops pathology in the late stages of PD and is expected to still have robust populations of neurons (unlike the substantia nigra, where neuron loss is severe at this disease stage).

**Table 1:** **Cohort characteristics** including disease status, age at death in years, postmortem interval (PMI) in hours, and race. One male was missing race data. One male control sample was removed during quality control, leaving n=99 in the final analyses. SD = standard deviation

| Variables | Male (n = 63) |  | Female (n = 37) |  |
| --- | --- | --- | --- | --- |
| | Mean ± SD<br>or N (%) | Range | Mean ± SD<br>or N (%) | Range |
| *Disease Status* |  |  |  |  |
| Control | 30 (47.6%) |  | 20 (54.1%) |  |
| Parkinson's Disease | 33 (52.4%) |  | 17 (45.9%) |  |
| *Age at Death* |  |  |  |  |
| Control | 79.1 ± 9.1 | 53-93 | 82.2 ± 13.1 | 52-95 |
| Parkinson's Disease | 79.2 ± 7.4 | 64-95 | 79.4 ± 5.5 | 70-87 |
| *PMI* |  |  |  |  |
| Control | 3.28 ± 0.82 | 2.16-5.5 | 3.05 ± 0.97 | 1.25-5 |
| Parkinson's Disease | 3.28 ± 0.86 | 1.83-5.23 | 3.19 ± 0.8 | 1.75-4.42 |
| *Race* |  |  |  |  |
| White | 62 (98.4%) |  | 37 (100%) |  |
| Unidentified | 1 (1.6%) |  | 0 (0%) |  |

### Validation of neuronal enrichment

To validate enrichment of NeuN^+^ nuclei via MACS prior to experimentation, we quantified total nuclei and NeuN^+^ nuclei for six test samples using flow cytometry. A representative quantification plot is included in **Figure 1a**. Plots for all six samples are available as a supplementary file (**Supplementary File 4**). For the test samples, the mean % NeuN^+^ was 92.0% (range = 89.2 – 95.2%). As a secondary validation, we estimated proportions of neuronal and glial cell types for all experimental MACS-sorted parietal cortex samples using the *CETS* package during EPIC array data processing. One sample from the male control group had an estimated neuronal population of only 0.03% in the *CETS* analysis. This sample was removed prior to all downstream differential methylation testing. The average proportion of neurons in the remaining 99 samples was estimated to be 83.8% by *CETS* analysis (**Figure 1b**).

**Figure 1:**
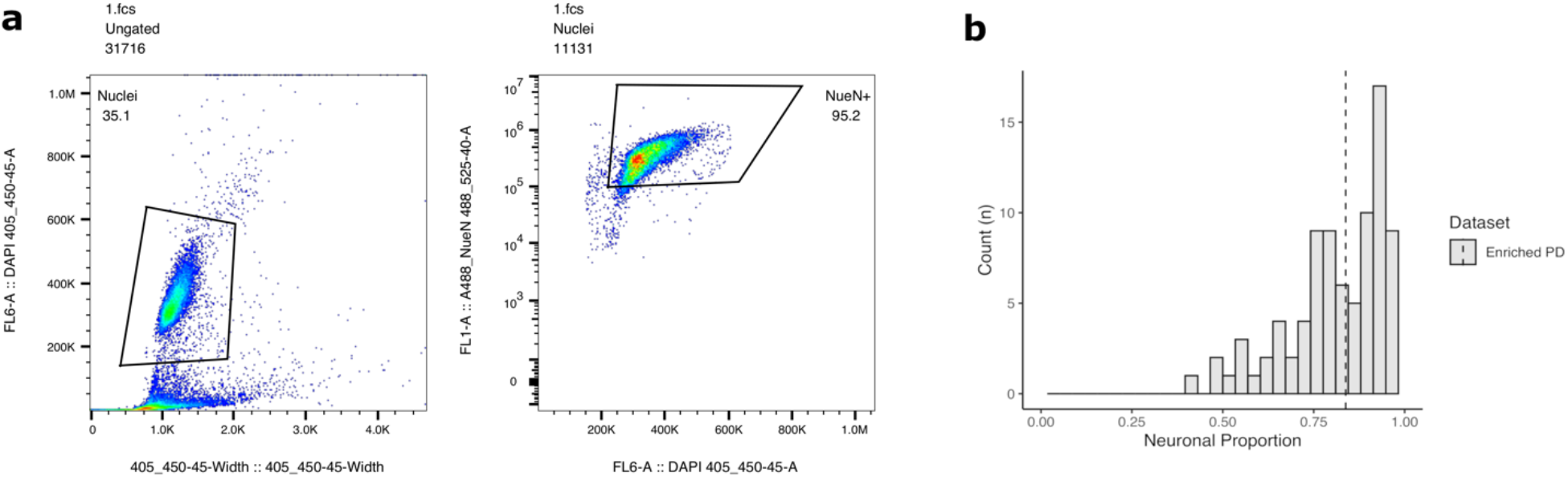
Neuronal enrichment validation by flow cytometry (a) and CETS analysis (b). a) NeuN^+^ nuclei were identified using DAPI pulse area vs. width using the 405-450/45 channel for all nuclei (left) followed by NeuN-AlexaFluor488 bright events 488-525/40 (right). b) Histogram of estimated neuronal proportion (vs. glial) for enriched PD NeuN^+^ nuclei population. Proportions of neuronal and glial cell types were estimated for MACS-sorted NeuN^+^ nuclei samples using the *CETS* R package during EPIC array data processing. Average proportion of neurons (vertical dotted line) was estimated to be 83.8% across all n=99 samples included in analysis.

### Differential testing for differentially methylated CpGs

In our sex-stratified models for differential 5-mC testing, we identified 3 differentially methylated CpGs (DMCs) in males and 87 significant DMCs in female parietal cortex associated with PD (p-value < 9×10^-8^) (**Supplementary Files 5 and 6**). These DMCs annotated to 3 unique gene IDs in the males and 85 unique gene IDs in the females. The sex-stratified DMC models included a sigma term to test for differences in mean by disease state while accounting for potential differences in variability. This p-value cutoff was selected based on recommendations in a recent study of epigenome-wide association studies.^48^ The sex-stratified DMC models included a sigma term to test for differences in mean by disease state while accounting for potential differences in variability. These results were sex-specific, and there was no overlap between any of the identified male and female DMCs by probe ID. The most significant DMC in males was in a CpG island within the 5’UTR of the *PARK7* (DJ-1) gene (p-value = 1.38×10^-9^). This *PARK7* DMC showed male-specific hypomethylation in PD cortical neurons compared to control (**Figure 2a**). In females, the most significant DMC was in the 5’UTR of the *ATXN1* gene (p-value = 1.84×10^-17^). This *ATXN1* DMC showed female-specific hypermethylation in PD cortical neurons compared to control (**Figure 2b**). We also identified a female-specific hypomethylated DMC within the gene body of the *SLC17A6* locus (p-value = 1.45×10^-10^) (**Figure 2c**).

**Figure 2:**
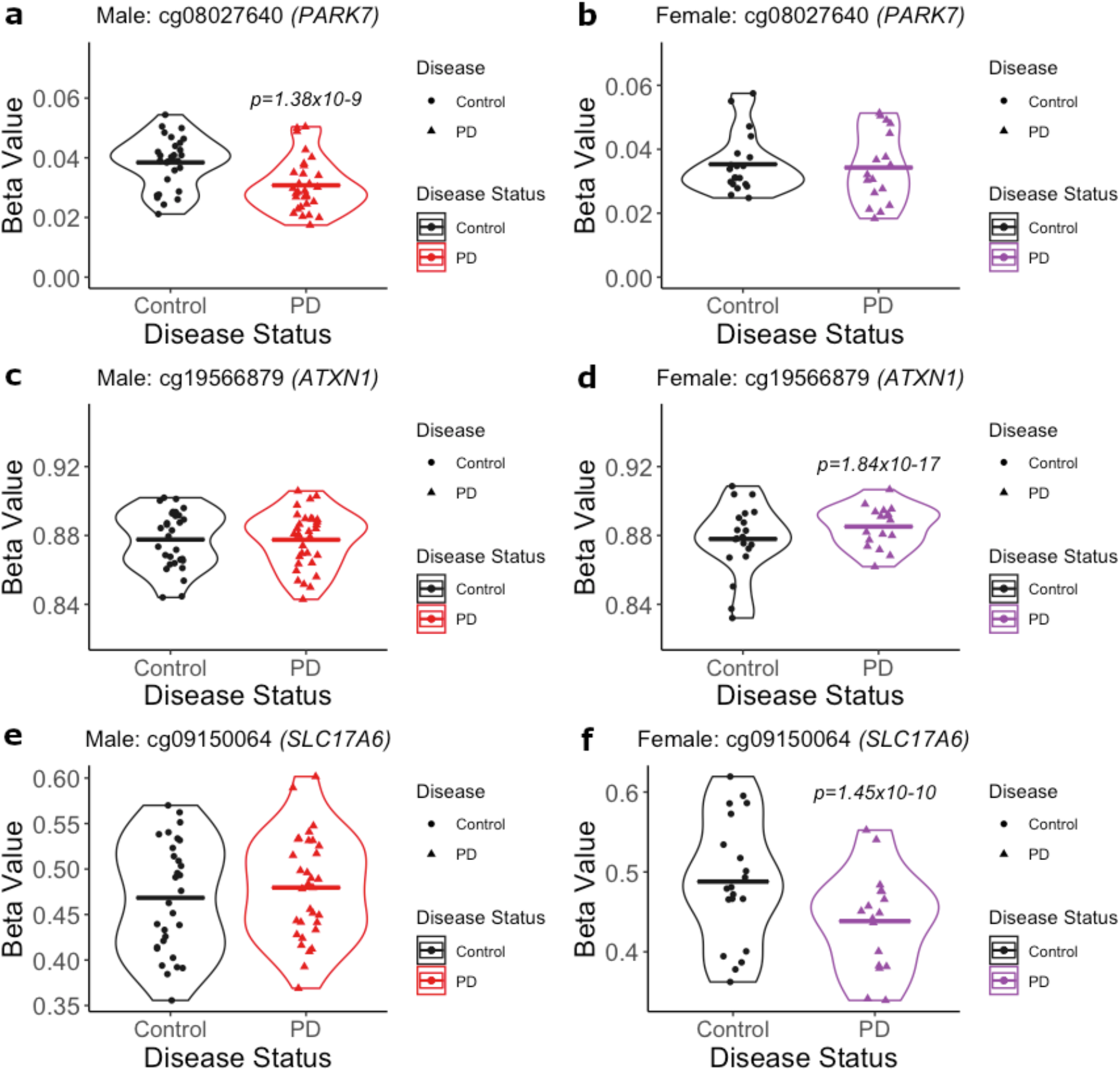
Dot plots of significant male and female DMCs by Parkinson’s disease. a) In males, the most significant DMC was annotated to the *PARK7* locus (cg08027640; p-value = 1.38×10^-9^). b) There was no significant change in DNA methylation at cg08027640 in females. c) There was no significant change in DNA methylation at cg19566879 in males. d) In females, the most significant DMC was annotated to the *ATXN1* locus (cg19566879; p-value = 1.84×10^-17^). e) There was no significant change in DNA methylation at cg09150064 in males. f) In females, a significant DMC was annotated to the *SLC17A6* locus (cg09150064; p-value = 1.45×10^-10^). P-values indicate significant change in PD compared to control (cutoff for significance: p-value < 9×10^-8^).

### Differential testing for differentially methylated regions

In our sex-stratified models for differential genomic region testing, we identified 258 significant differentially methylated regions (DMRs) in males and 214 significant DMRs in female parietal cortex by Parkinson’s disease (minimum smoothed FDR < 0.05) (**Supplementary Files 7 and 8**). These DMRs annotated to 205 unique gene IDs in the males and 157 unique gene IDs in the females. Comparing these two lists of DMR by chromosomal location, 5 regions showed at least partial overlap between the two sexes (Error! Reference source not found.), and one region – annotated to the *PTPRN2* gene – showed exact, complete overlap in both males and females (**Figure 3a, 3b**). This region of complete overlap (chr7: 158093198 – 158093277) includes 3 CpGs – cg11293572, cg09992350, and cg27014435 – and is annotated to an intronic region within the *PTPRN2* gene body. Although this region was a DMR in both males and females, it was hypermethylated in male brains and hypomethylated in female brains. (Error! Reference source not found.; **Figure 3**). The other 4 regions of partial overlap – annotated to the *ZIC1, GALNT15, KIAA0040*, and *GFPT2* genes – also showed opposite relationships between PD status and DNA methylation in males and females (Error! Reference source not found.). In addition to overlapping DMRs, we also identified a female-specific DMR annotated to the *SLC17A6* gene, which included the DMC identified at this gene(**Figure 3d**). This DMR (chr11:22362708-22364961) includes 8 CpGs – cg21518089, cg04583232, cg24454829, cg17298751, cg09150064, cg24151995, cg20809470, and cg22371972 – and spans intron 1, exon 2, intron 2, and exon 2 of the *SLC17A6* gene. Lastly, we also identified a male-specific DMR annotated to the *NR4A2* gene (**Figure 3e**). This DMR (chr2:157186666-157186681) includes 2 CpGs – cg21226516 and cg11209121 – that are located at an exon-intron boundary in the gene body of *NR4A2*. These results underscore that PD-related changes in cortical neuron DNA modifications are sex-specific.

**Figure 3:**
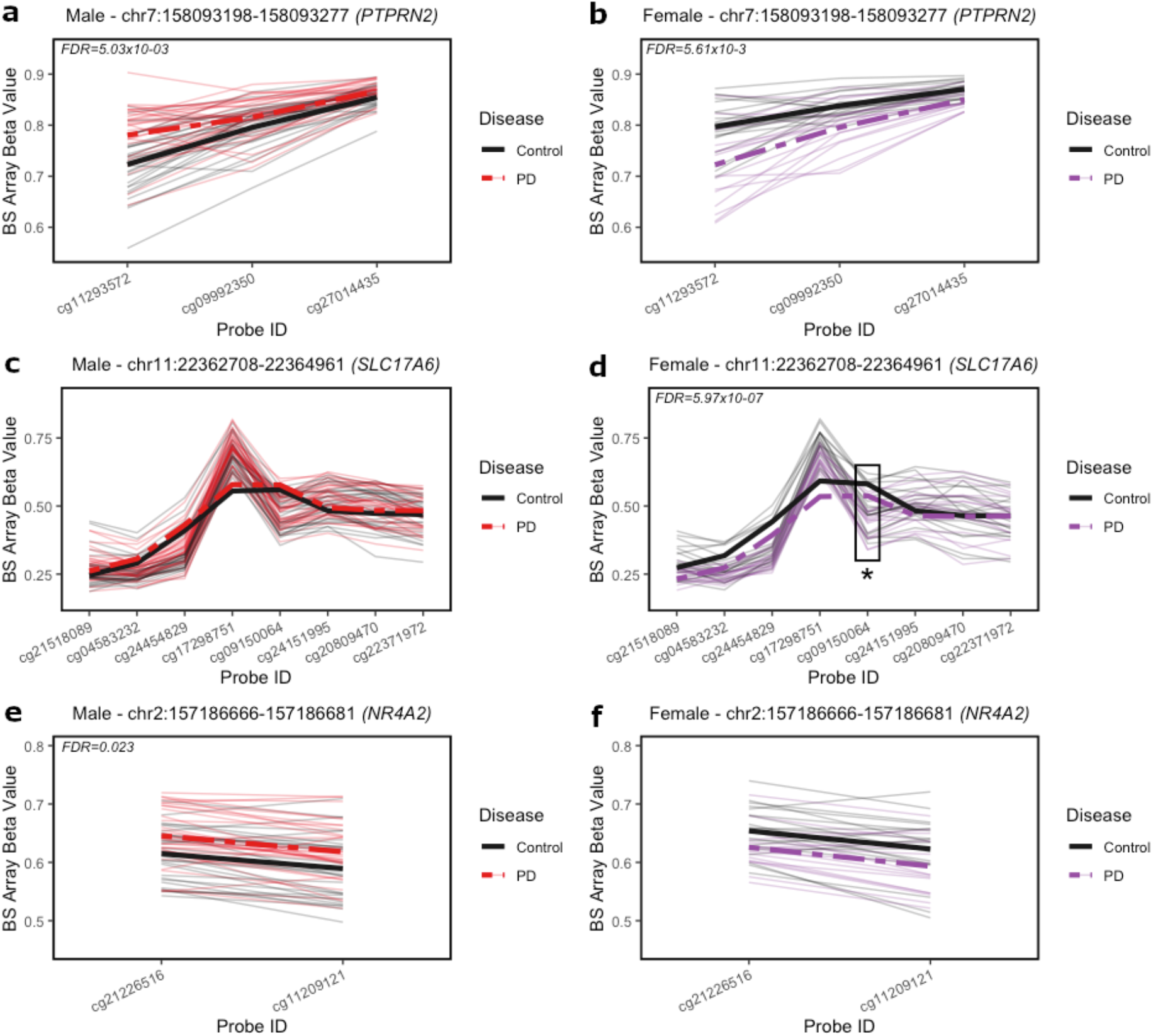
Spaghetti plots of *PTPRN2, SLC17A6*, and *NR4A2* DMRs in males and females. Beta values (y-axis) represent DNA methylation across CpGs included in DMRs annotated to *PTPRN2* (chr7: 158093198 – 158093277), *SLC17A6* (chr11:22362708-22364961), and *NR4A2* (chr2:157186666-157186681). a) In males, PD brains (red) exhibited significant hypermethylation at *PTPRN2* compared to control (black). b) In females, PD brains (purple) exhibited significant hypomethylation at *PTPRN2* compared to control (black). c) In males, PD brains (red) did not exhibit significant differential methylation at *SLC7A6* compared to control (black). d) In females, PD brains (purple) exhibited significant hypomethylation at *SLC17A6* compared to control (black). Significant DMC (cg09150064) indicated with box and asterisk. e) In males, PD brains (red) exhibited significant hypermethylation at *NR4A2* compared to control (black). f) In females, PD brains (purple) did not exhibit significant differential methylation compared to control (black). Significant DMRs are indicated with minimum smoothed FDR values < 0.05. Thick solid and dashed lines represent smoothed means by group across DMRs.

**Table 2:**
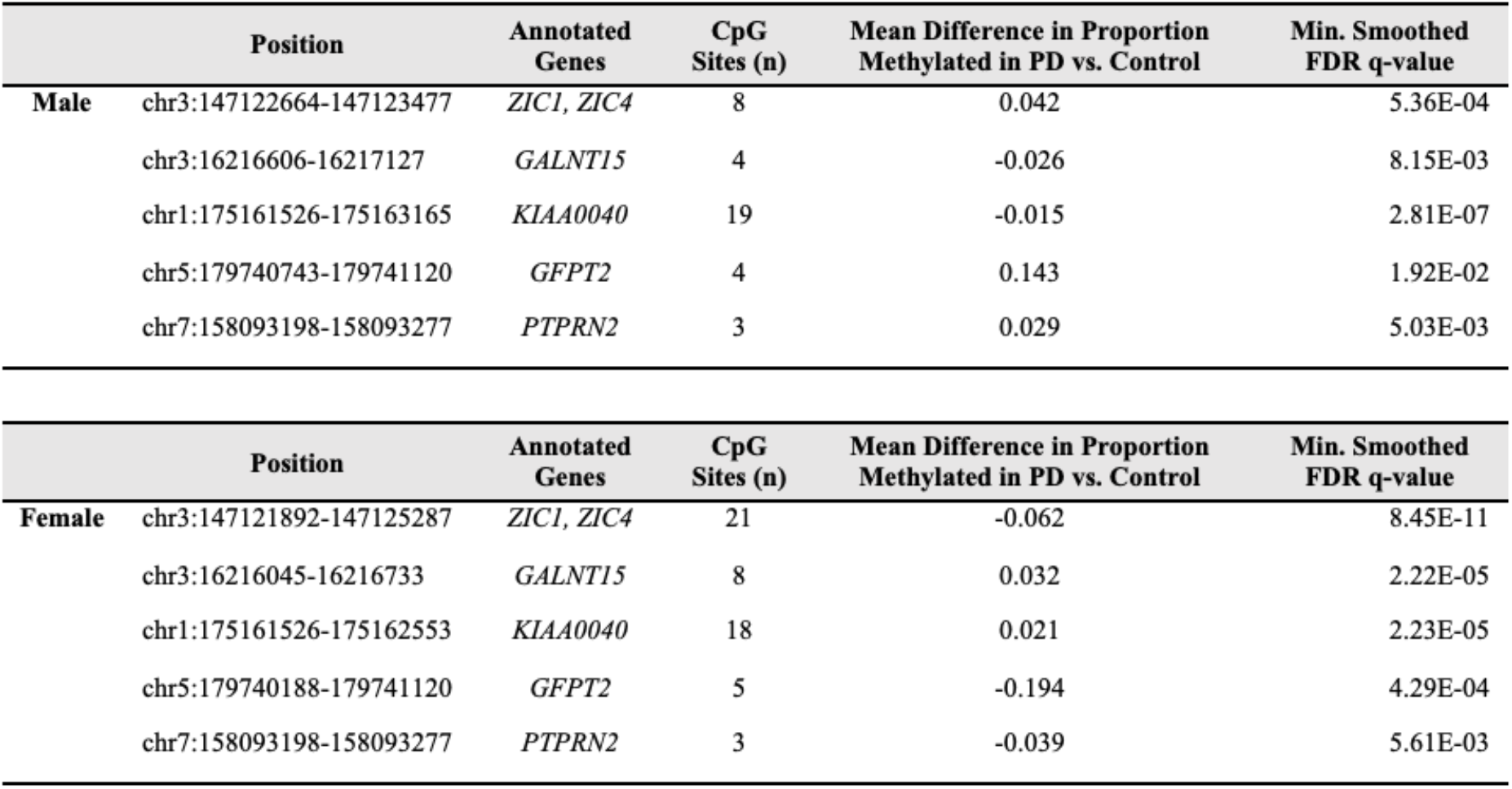
Differentially methylated regions (DMRs) by Parkinson’s Disease. Five significant DMRs (minimum smoothed FDR < 0.05) showed at least partial overlap in both sexes, as shown in the sex-stratified tables (Top = males; bottom = females). One region – annotated to the *PTPRN2* gene – showed exact, complete overlap in both males and females. All regions of overlap showed sex-specific directions of differential DNA methylation with PD. DMR modeling by Parkinson’s disease status included age and estimated glial cell proportion as covariates. FDR = false discovery rate.

### Gene ontology pathway enrichment

The *clusterProfiler* R package was used to determine whether DMCs annotated to genes enriched for specific gene ontology biological process (GOBP) pathways. GOBP pathway enrichment analysis was not performed on male DMCs due to the low number of significant CpGs (n=3) but was performed on the genes annotated to female hypomethylated DMCs and hypermethylated DMCs separately. Hypo- and hypermethylated DMCs were considered separately due to expected differences in associations between increased or decreased DNA methylation and gene regulation. Female hypermethylated DMCs did not show enrichment for any pathways, but female hypomethylated DMCs showed enrichment for 10 pathways, including several developmental pathways, neurotransmitter transport, neurotransmitter secretion, and signal release from synapse (**Figure 4a**). Several differentially methylated genes were included in these GO term pathways, including *SLC17A6* and *PTPRN2* (**Figure 4b**), the latter of which showed differential DNA methylation at the region level in both males and females.

**Figure 4:**
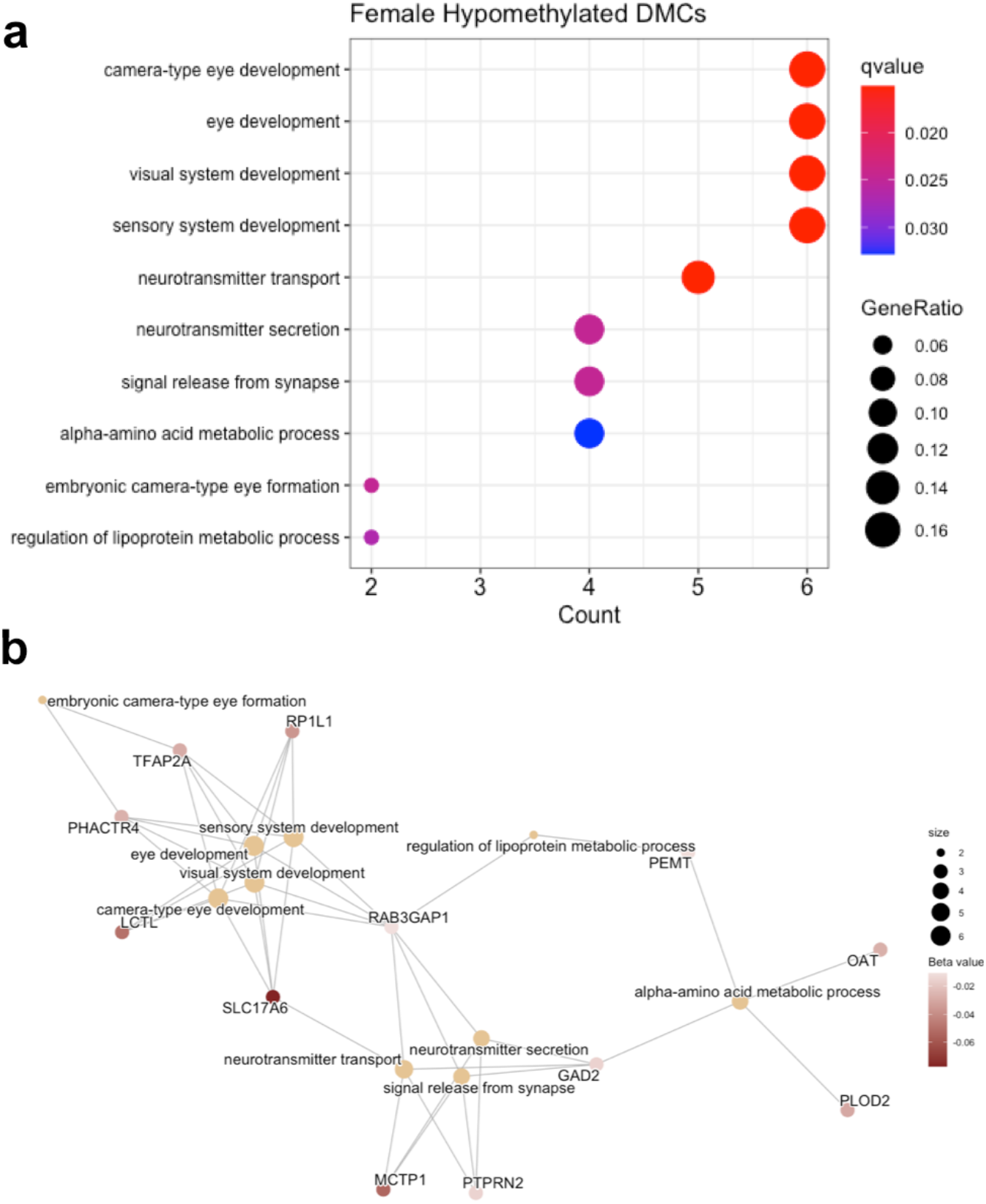
GO Term enrichment dot plot and gene-concept network plot for female hypomethylated DMCs. a) Female hypomethylated DMCs showed enrichment for 10 pathways, including neurotransmitter transport, neurotransmitter secretion, and signal release from synapse. The x-axis is the number of genes in each pathway, color is used to represent FDR q-value (qvalue), and size is used to represent GeneRatio, the ratio of differentially methylated genes in each GO term to the total number of genes input into the hypergeometric test. b) Several differentially methylated genes were included in the enriched pathways, including *PTPRN2*, which also showed differential DNA methylation at the region level in both males and females. The size of each node is used to represent the number of genes in each GO term, and color represents the magnitude of decrease in DNA methylation (beta value) for each annotated probe. Connections between genes and GO terms represent inclusion of the gene in that GO term.

The *clusterProfiler* R package was also used to determine whether DMRs annotated to genes enriched for specific gene ontology biological process (GOBP) pathways. In both males and females, hypermethylated DMRs did not show any enrichment for GOBP terms. Meanwhile, for the hypomethylated DMRs, females showed significant enrichment for 6 GOBP terms (q-value < 0.05), including cell fate commitment and glucose homeostasis (**Supplementary File 9**). These enriched pathways for the female hypomethylated DMRs are collectively related to cell fate and metabolism. In male hypomethylated DMRs, 4 pathways approached significance (q-value = 0.06), including semaphorin-plexin signaling pathways involved in axon guidance and neuron projection guidance (**Supplementary File 9**).

### Epigenetic clock analysis

In our epigenetic clock analysis, estimated epigenetic age significantly predicted chronological age in both sexes, with estimated age (“EpiAge”) significantly associated with chronological age at death (males: Beta coefficient = 1.1929, p-value = 8.97×10^-9^; females: Beta coefficient = 1.3515, p-value = 2.51×10^-9^). These results confirm that this is a well-calibrated epigenetic clock where there was an approximate 1 unit increase in estimated epigenetic age with a 1 unit increase in chronological age (**Table 3**). However, in our data, PD status did not modify trajectories of epigenetic aging in either sex (**Figure 5**, **Table 3**).

**Figure 5:**
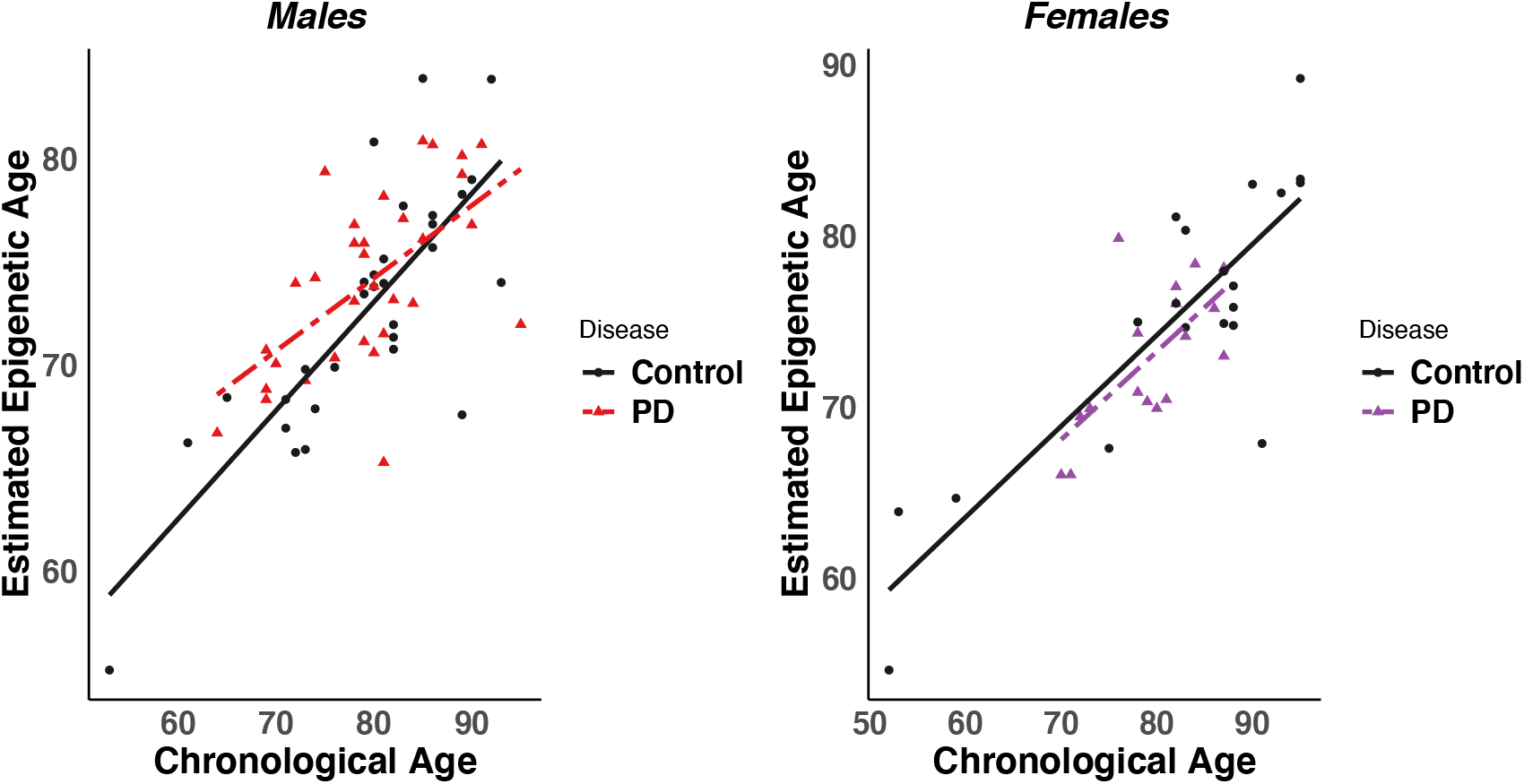
Scatterplots of male and female estimated epigenetic age vs. chronological age at death. Estimated epigenetic age showed a significant positive association with chronological age in males (left) and females (right). However, PD status did not significantly modify trajectories of epigenetic age. Control samples are shown as black dots, and PD samples are shown as red or purple triangles for males and females, respectively.

**Table 3:** Epigenetic clock linear regression results. Models were run for male subjects (top), and female subjects (bottom). EpiAge refers to estimated epigenetic age. DiseasePD refers to the PD status variable (reference group = control). EpiAge:DiseasePD is the interaction term between estimated epigenetic age and PD status that determines whether PD brains exhibit altered trajectories of epigenetic age compared to control. Estimate = beta coefficients from regression model; Std. Error = standard error.

| Male (n=62) |  |  |  |  |
| --- | --- | --- | --- | --- |
| Coefficients: | Estimate | Std. Error | t-value | p-value |
| EpiAge | 1.1929 | 0.1778 | 6.711 | 8.97x10<sup>-9</sup> |
| Disease PD | 11.8959 | 22.137 | 0.537 | 0.593 |
| EpiAge:DiseasePD | -0.1788 | 0.3004 | -0.595 | 0.554 |

**Table 3: Epigenetic clock linear regression results.**
| Female (n=37) |  |  |  |  |
| --- | --- | --- | --- | --- |
| Coefficients: | Estimate | Std. Error | t-value | p-value |
| EpiAge | 1.3515 | 0.1672 | 8.081 | 2.51x10<sup>-9</sup> |
| Disease PD | 35.7668 | 28.8822 | 1.238 | 0.224 |
| EpiAge:DiseasePD | -0.484 | 0.3927 | -1.233 | 0.226 |

## Discussion

Here, we report the first genome-wide analysis of DNA methylation in an enriched neuronal population from PD post-mortem parietal cortex. We identified sex-specific PD-associated methylation changes in a total of 434 unique genes, including *PARK7* (DJ-1), *SLC17A6* (VGLUT2), *PTPRN2* (IA-2β), *NR4A2* (NURR-1), and other genes involved in developmental pathways, neurotransmitter packaging and release, and axon and neuron projection guidance (**Figure 2, Figure 3, Figure 4**). Our data did not show accelerated epigenetic aging in PD (**Figure 5, Table 4**). This study expands on the existing literature through the utilization of cell-sorted cortical neurons, inclusion of both sexes, and estimation of epigenetic age.

### Sex-specificity of PD-associated differential DNA methylation

While we expected to identify differences in DNA methylation in male and female samples due to the known sex differences in susceptibility and progression of PD,^29–31^ the lack of overlap between sexes was striking. All the identified DMCs and almost all DMRs were specific to either males or females. Furthermore, of the 5 DMRs that showed either complete or partial overlap between sexes, all 5 showed opposite directions of change in male and female samples (Error! Reference source not found.). As this is the first genome-wide study of DNA methylation in PD that analyzed male and female samples separately, this finding underscores the importance of stratifying data by sex rather than including as a covariate during analysis. Inclusion of a sex covariate can correct for changes in baseline DNA methylation level. However, a simple covariate will not capture changes in direction of effect between sexes, as were seen at all five DMRs with overlap between males and females (Error! Reference source not found.). Instead, a sex:disease interaction term must be included, but this leads to increased sample size requirements, especially in human cohorts with complex, multivariate models. To remove the requirement for an interaction term and preserve statistical power, we elected to stratify our analyses by sex. Our identified differences in DNA methylation between male and female subjects highlight the need for future PD epigenome-wide studies to stratify by sex and provide insight into potential mechanisms underlying the well-recognized sex differences in PD susceptibility and progression.

### Male-specific PD-associated DNA hypomethylation at the PARK7 locus

The most significant DMC in male subjects was in a promoter-associated CpG island within the 5’-UTR of the *PARK7* locus, which showed PD-related hypomethylation (**Figure 2**). Mutations in the *PARK7* locus, which encodes the DJ-1 protein, cause autosomal recessive, early-onset Parkinson’s disease and oxidized DJ-1 has been observed in brains of idiopathic PD patients.^49,50^ A review of DJ-1 genetics identified a PD-linked variant in the DJ-1 5’UTR, supporting that epigenetic regulation at this genomic region could play a role in idiopathic disease.^50^ DJ-1 has multiple known neuroprotective functions – transcriptional regulator, molecular chaperone, and antioxidant – and is thought to play a role in both familial and idiopathic forms of PD.^51–54^ While DJ-1 has been studied extensively in the context of PD, ours is the first study to identify an association between DNA methylation at the *PARK7* locus and PD. Two previous studies did not identify significant effects of PD status on DNA methylation at the *PARK7* locus in blood leukocytes or post-mortem brain tissue.^55,56^ However, both studies used targeted bisulfite pyrosequencing and looked at different sites within the *PARK7* locus. In addition, one study was performed in blood leukocytes and may may not reflect DNA methylation in brain tissue.^55^ The other used human post-mortem brain tissue, but had a very small sample size and used bulk tissue samples.^56^ Their results do not contradict our *PARK7* results and suggest that follow up studies should examine the entire locus. In addition, these differences in study design emphasize the importance of considering tissue choice, genomic location, sample size, and cell type when interpreting DNA methylation data. While we did not assess RNA or protein levels of DJ-1 in this study, our data suggest that epigenetic regulation of transcription of this locus may play a role in idiopathic PD.

### Female-specific PD-associated DNA hypermethylation at the ATXN1 locus

The most significant DMC in female subjects was located within the 5’UTR of the *ATXN1* gene, which showed PD-related hypermethylation (**Figure 2**). Based on the Illumina EPIC manifest, this CpG does not overlap any known transcription factor binding sites, but it does fall within a DNase hypersensitivity region (chr6:16333840-16334215). Thus, the DNA region including this CpG is accessible for cleavage by DNase enzymes, which often occurs at regulatory regions.^57^ *ATXN1* encodes Ataxin-1 (ATXN1), which is involved in transcriptional repression of a large number of target genes and is necessary for normal brain development and function.^58,59^ Expanded polyglutamine repeats in this gene cause spinocerebellar ataxia type 1 (SCA1).^58,60^ Data show that dysfunction of this protein in cortical neurons disrupts neurodevelopment and causes a spectrum of neurobehavioral phenotypes (Lu 2017 – 69). Although this gene has not been previously studied specifically in the context of PD, this new evidence showing altered epigenetic regulation of the *ATXN1* gene in PD brains opens new avenues of research. It is possible that epigenetic regulation at this locus leads to changes in expression of ATXN1 and its target genes that affect the susceptibility of cortical neurons to PD pathology.

### Male-specific PD-associated changes in DNA methylation at NR4A2

In males, we also identified PD-related hypermethylation in an exon just downstream of an exon-intron boundary in the gene body of *NR4A2* (**Figure 3**). *NR4A2* encodes the nuclear receptor related-1 (NURR1) protein. NURR1 is a transcription factor critical for dopaminergic neuron development and maintenance. Previous research suggests that altered regulation of the *NR4A2* gene may contribute to PD pathogenesis.^61–63^ In a previous animal model study, we showed that developmental exposure to the organochlorine pesticide dieldrin, which is known to be associated with increased PD risk, led to sex-specific hypermethylation in the *Nr4a2* gene body in female mice.^64^ These previous results, along with our present data, suggest that sexspecific epigenetic regulation at the *NR4A2* gene may play a role in idiopathic PD risk.

### Female-specific PD-associated DNA hypomethylation at the SLC17A6 locus

In females, we also identified PD-related hypomethylation in an intron within the *SLC17A6* gene body (**Figure 2, Figure 3**). *SLC17A6* encodes the vesicular glutamate transporter 2 (VGLUT2), which has been implicated in PD pathogenesis by multiple lines of evidence.^65^ Studies in *in vivo* and *in vitro* models show increased VGLUT2 expression in dopaminergic (DA) neurons in response to the neurotoxicants, rotenone, 6-OHDA and MPP^+^.^66,67^ Deletion of VGLUT2 in mice exacerbates the neurotoxic effects of 1-methyl-4-phenyl-1,2,3,6-tetrahydropyridine (MPTP) exposure.^68^ Many studies suggest that neurons that co-express VGLUT2 and VMAT2 show differentially vulnerability in PD and differences in neuronal structure.^69,70^ Our study did not assess DNA methylation in DA neurons of the nigrostriatal pathway, which have been the subject of much of this work on VGLUT2 in the context of PD. However, the observed change in DNA methylation within the *SLC17A6* gene in parietal cortex raises questions about an additional role for VGLUT2 in cortical neurons in PD.

### Sex-specific PD-associated changes in DNA methylation at PTPRN2

In our regional analysis, one DMR – annotated to the *PTPRN2* gene – showed exact, complete overlap in both males and females (Error! Reference source not found.). Although the identified *PTPRN2* DMR was differentially methylated in both males and females, it was hypermethylated in brains from male PD patients and hypomethylated in brains from female PD patients (**Figure 3**). Altered methylation of this gene has been previously implicated in PD in a study of longitudinal blood samples that showed an association between hypomethylation of a different CpG within the *PTPRN2* gene in whole blood and faster motor progression in PD.^71^ *PTPRN2* encodes Protein Tyrosine Phosphatase Receptor Type N2 (IA-2β), which is expressed on dense core and synaptic vesicles. Deletion of this gene and the related IA-2 protein in mice leads to reduced levels of norepinephrine, dopamine and serotonin in the brain and decreased release of DA, GABA, and glutamate from synaptosomes.^72^ In addition, existing data implicates this gene in PD. One previous study found decreased expression of *PTPRN2* in the SN of PD patients, while another found increased expression in DA neurons from PD patients with *LRRK2* G2019S mutations.^73,74^ Interestingly, given the known role of environmental exposures in PD and work in our lab investigating pesticide exposures and PD, ^64,75^ another study found an association between pesticide exposure and hypomethylation at a CpG in *PTPRN2* in blood.^76^ Taken together, our data adds to a growing collection of evidence supporting a role for epigenetic regulation of *PTRPN2* in PD.

### Sex-specific pathways enriched for DNA methylation changes

Beyond our gene-level differences, pathway analysis of differentially methylated genes implicates distinct pathways in male and female subjects (**Figure 4**). Female-specific differentially methylated genes were enriched for pathways including neurotransmitter transport, neurotransmitter secretion, and signal release from synapse (**Figure 4c**). These pathways were not enriched in differentially methylated genes identified in male subjects. Instead, the semaphorin-plexin signaling pathways involved in axon guidance and neuron projection guidance were enriched in male-specific differentially methylated genes. These pathways included *PLXNB1, PLXNC1*, and *PLXNB3*, three genes that encode proteins in the plexin family, which act as transmembrane receptors for the semaphorins. The semaphorin-plexin signaling pathway is a key regulator of morphology and motility in many different cell types, including those that make up the nervous system, and is known to play a role in axon guidance.^77,78^

### PD is not associated with accelerated epigenetic aging

Over the past decade, recognition has been growing that the epigenome, especially DNA methylation, can be used to closely predict chronological age.^41,79–81^ Based on this, researchers established epigenetic clocks that use loci with the most predictable DNA methylation levels to provide estimates of epigenetic (or biological) age.^79^ In a previous study using one of these epigenetic clocks, researchers found accelerated epigenetic aging in PD blood samples compared to control.^45^ However, to our knowledge, no studies have tested whether epigenetic aging is accelerated in PD brain tissue. Here, we used an epigenetic clock specifically designed for human cortical tissue to estimate epigenetic age of control and PD parietal cortex samples.^82^ In our analyses, we showed that our epigenetic clock was well calibrated to predict chronological age at death, but found no evidence of significant epigenetic age acceleration by PD status in our cohort (**Figure 5**). These data suggest that accelerated epigenetic aging may not play a role in PD development, but this result should be further investigated in a larger cohort.

### Conclusions

Our study marks the first genome-wide analysis of DNA methylation in an enriched neuronal population from human brain tissue. However, there are some limitations to our current analysis that create opportunities for future studies. Our results do not distinguish between DNA methylation and DNA hydroxymethylation.^34^ These marks show distinct genomic distributions and associations with gene expression, particularly in the brain, as well as differential responses to environmental exposures.^9,83,84^ As such, DNA methylation and DNA hydroxymethylation may play discrete roles in neurodegenerative diseases.^9,84,85^ It is critical for future studies to use modifications to the bisulfite conversion protocol, like oxidative BS, or enzymatic conversion methods that can distinguish between these marks. In addition, because we used the Illumina EPIC array in this study, we did not achieve a true “genome-wide” wide analysis. This array is not specifically designed for brain-related disease, and many genomic regions of interest to the PD field are not covered on the array. True genome-wide analysis requires whole-genome bisulfite sequencing, but it remains cost-prohibitive to scale up for a large cohort. Our study had a relatively small sample size, especially when stratified by sex. Based on the recommendations published by Mansell et al., a sample size of at least N=200 (n=100 per disease group) per sex would be more appropriate.^48^ Future studies should include larger cohorts or be combined with with public data in the field. Finally, we only analyzed the neuron-enriched population from our MACS sorted samples. Given the growing body of evidence supporting the important of various non-neuronal cell types in PD, carrying out a similar analyses in these neuron-depleted samples is also important to being to understand the role of DNA methylation in other cell types. Despite the limitations of our study, this is the first genome-wide analysis of DNA methylation in enriched neurons from human postmortem brain tissue. We identified sex-specific changes in DNA methylation at genes known to be related to PD as well as genes not previously implicated in PD. Together, our data support the growing recognition that epigenetic regulation is an important mechanism in PD pathogenesis and point to specific genes and pathways for further study.

## Methods

### Magnetic-activated cell sorting

NeuN-positive (NeuN^+^) nuclei were enriched from 100 mg of flash frozen parietal cortex tissue using a two-stage magnetic-assisted cell sorting (MACS) method (**Figure 1**). First, 100 mg of frozen tissue were briefly thawed on ice and homogenized in a 2 mL, 1.4 mm ceramic bead tube (Thermo Fisher Scientific, Cat. # 15-340-153) with 1 mL of Nuclear Extraction Buffer (NEB) for 10 sec at 4 m/s. NEB consisted of 0.32M sucrose, 0.01M Tris-HCl pH 8.0, 0.005M CaCl_2_, 0.003M MgCl_2_, 0.0001M EDTA, and 0.1% Triton X-100, up to a stock volume of 1 L using water. Immediately prior to use, 0.001M DTT was added to NEB. Homogenized samples were loaded into a 13 mL ultracentrifuge tube (BeckmanCoulter, Cat. # 331372) with 4 mL of NEB. Using a glass pipette, 7 mL of sucrose solution was pipetted down the side of each sample tube to create a sucrose gradient. Sucrose solution consisted of 1.8M sucrose, 0.01M Tris-HCl pH 8.0, 0.003M MgCl_2_, up to a stock volume of 1 L using water. Immediately prior to use, 0.001M DTT was added to NEB. After addition of sucrose, samples were spun at 4°C, 24,000 rpm in the Sorvall Wx+ Ultracentrifuge in a swing bucket rotor (TH-641). Once the centrifugation was complete, the supernatant and debris layer found at the concentration gradient were both removed with the use of a vacuum, while being careful not to disturb the pellet containing the nuclei at the bottom of the tube. Next, 1 mL of primary antibody (anti Neun 488 – Millipore, Cat. # MAB377X) in MACS buffer was added to each nuclei pellet and placed on ice for 10 minutes. MACS buffer consisted of 0.5% Bovine Serum Albumin solution (Sigma-Aldrich, Cat. # A1595) in PBS pH 7.2 (Gibco, Cat. # 20012-027). Samples were then mechanically pipetted up and down 10-15 times to completely dissolve the nuclei pellet within the primary antibody-MACS buffer solution. This solution of nuclei was then transferred to a 2 mL tube and incubated for 60 minutes at 4°C. After incubation, 40 μL of MACS Microbeads (anti-mouse IgG Microbeads – Miltenyi, Cat. # 130-048-401) were added to each sample. Samples were then inverted 4-5 times and incubated at 4°C for 30 minutes. After incubation, nuclei were centrifuged at 300 x g for 10 minutes. Supernatant was then removed, and the nuclei were resuspended in 2 mL of MACS buffer and transferred to a MACS MS column (MS Separation columns – Miltenyi, Cat. # 130-042-201) that was pre-washed with MACS buffer and attached to the Miltyeni OctoMACS™ Separator. Positive selection of NeuN^+^ cells was then performed according to the standard MACS MS Columns protocol available from Miltenyi Biotec. After the first round of magnetic separation, NeuN^+^ nuclei were run through a separate, second MACS MS column to maximize cell type enrichment (**Figure 6**).

**Figure 6:**
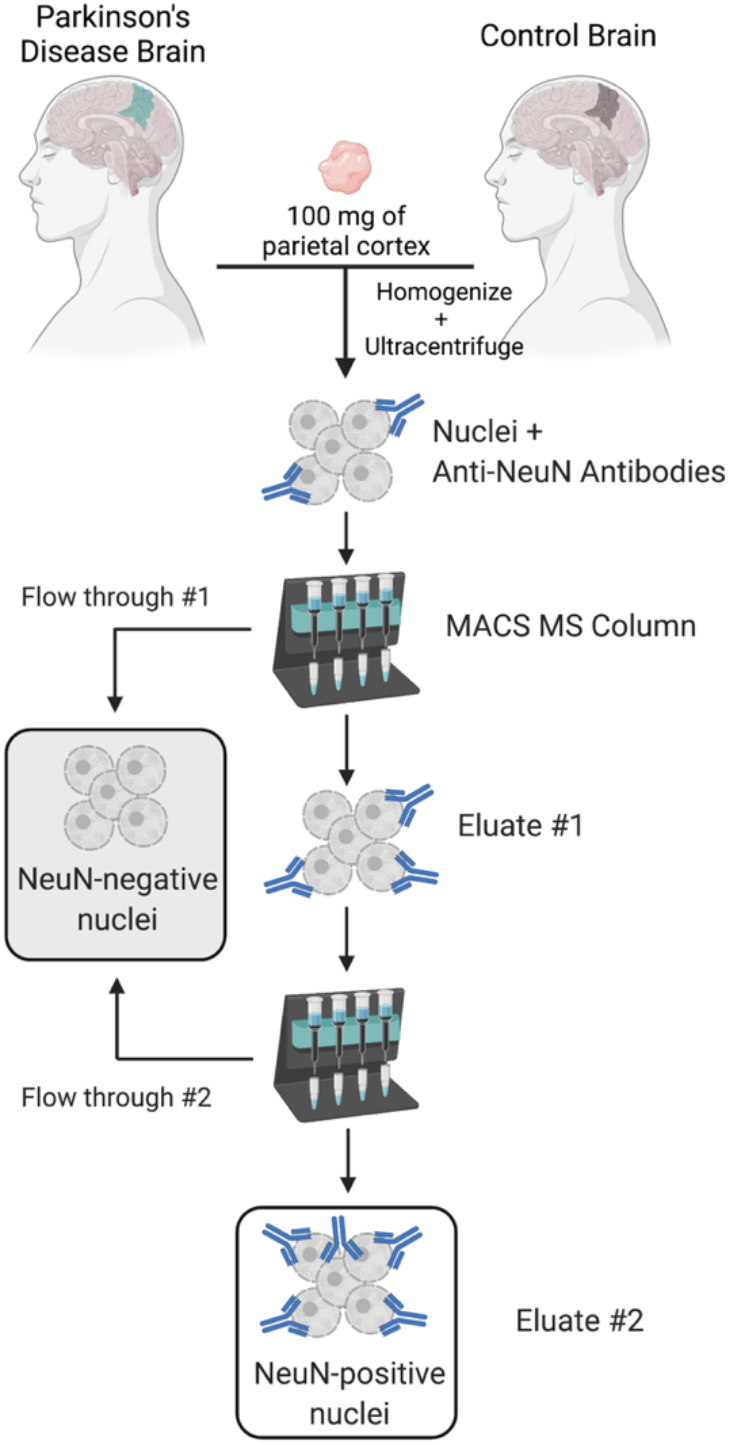
NeuN^+^ nuclei enrichment workflow. Nuclei were enriched from 100 mg of flash frozen parietal cortex tissue using a two-stage magnetic-activated cell sorting (MACS) method. After the first round of magnetic separation, NeuN^+^ nuclei were run through a separate, second column to maximize cell-type enrichment in final eluate.

### Enrichment validation using flow cytometry

Isolated nuclei were analyzed for flow cytometry on a CytoFlex S (Beckman Coulter), and data were analyzed using FlowJo V10. NeuN^+^ nuclei were identified using DAPI (pulse area vs. width using the 405-450/45 channel for all nuclei) followed by NeuN-AlexaFluor488 bright events (488-525/40).

### DNA extraction, bisulfite treatment, and EPIC arrays

DNA was isolated from enriched NeuN^+^ nuclei using the Qiagen QIAamp DNA Micro Kit (Cat. # 56304), with some modifications to maximize yield. Given that samples were already homogenized during nuclei isolation, the sample lysis and incubation steps of the QIAamp DNA Micro Kit protocol were removed. Instead, 20 uL of proteinase K were added directly to each MACS eluate. Samples were then vortexed for 15 seconds and incubated at room temperature for 15 minutes. In addition, the optional carrier RNA was added to Buffer AL, the incubation time after addition of 100% ethanol was increased to 10 minutes, the incubation time for the elution buffer was increased to 5 minutes, and the final elution step was repeated using 10 mM Tris-HCl pH 8.0.

Intact genomic DNA yield was quantified by Qubit fluorometry (Life Technologies). Bisulfite conversion was performed on 500 ng genomic DNA using the TrueMethyl Array kit (Cambridge Epigenetix). All conversion reactions were cleaned using SPRI-bead purification and eluted in Tris buffer. Following elution, BS-converted DNA was denatured and processed through the EPIC array protocol. The EPIC array contains ~850,000 probes that query DNA methylation at CpG sites across a variety of genomic features, including CpG islands, RefSeq genic regions, ENCODE open chromatin, ENCODE transcription factor binding sites, and FANTOM5 enhancer regions. To perform the assay, converted DNA was denatured with 0.4 N sodium hydroxide. Denatured DNA was then amplified, hybridized to the EPIC bead chip, and an extension reaction was performed using fluorophore-labeled nucleotides per the manufacturers protocol. Array BeadChips were scanned on the Illumina iScan platform.

### EPIC array data processing

After scanning on the iScan platform, BeadChip IDAT files were imported into R and processed using an in-house bioinformatics pipeline (**Figure 7**). This pipeline combined the *minfi* (version 1.22.1), *ChAMP* (version 2.14.0), *posibatch* (version 1.0), *ENmix* (version 1.12.4), *CETS, ssNoob*, and *dplyr* packages in R. Quality control was assessed for internal control probes using the *ENmix* plotCtrl function. Failed probes were identified as those with a detection p-value > 0.01 in any sample. Failed probes were then removed from downstream analyses when detection p-value was > 0.01 in > 5% of samples. No samples exceeded the selected threshold of failed probes (>10%) to warrant removal prior to analysis. Cross-reactive probes and probes containing SNPs were removed based upon previous identification.^86,87^ After removal of technical artifacts, dye-bias correction was performed with *ssNoob* within *minfi*^88^ The proportion of neuronal vs. glial cells in each sample was estimated with *CETS*.^89^ Samples with estimated glial cell to neuronal cell proportion > 0.90 were removed from analysis. This removed one sample from the male data set, leaving n=62 males. Batch effects were assessed using the *ChAMP* package,^90^ and then a beta value matrix for all samples was corrected for batch and positional effects using the*posibatch* R package.^91^ After beta value estimation, we filtered out samples with mean 5-mC beta value < 0.01. This beta value cutoff was instituted due to increased variability and decreased interpretability of beta values at such low levels. As a final step, we removed probes that were not annotated to a gene in the Illumina EPIC array manifest to ensure interpretability of results after modeling. After all filtering steps, 552,332 probes remained for differential DNA methylation testing in the sex-stratified datasets.

**Figure 7:**
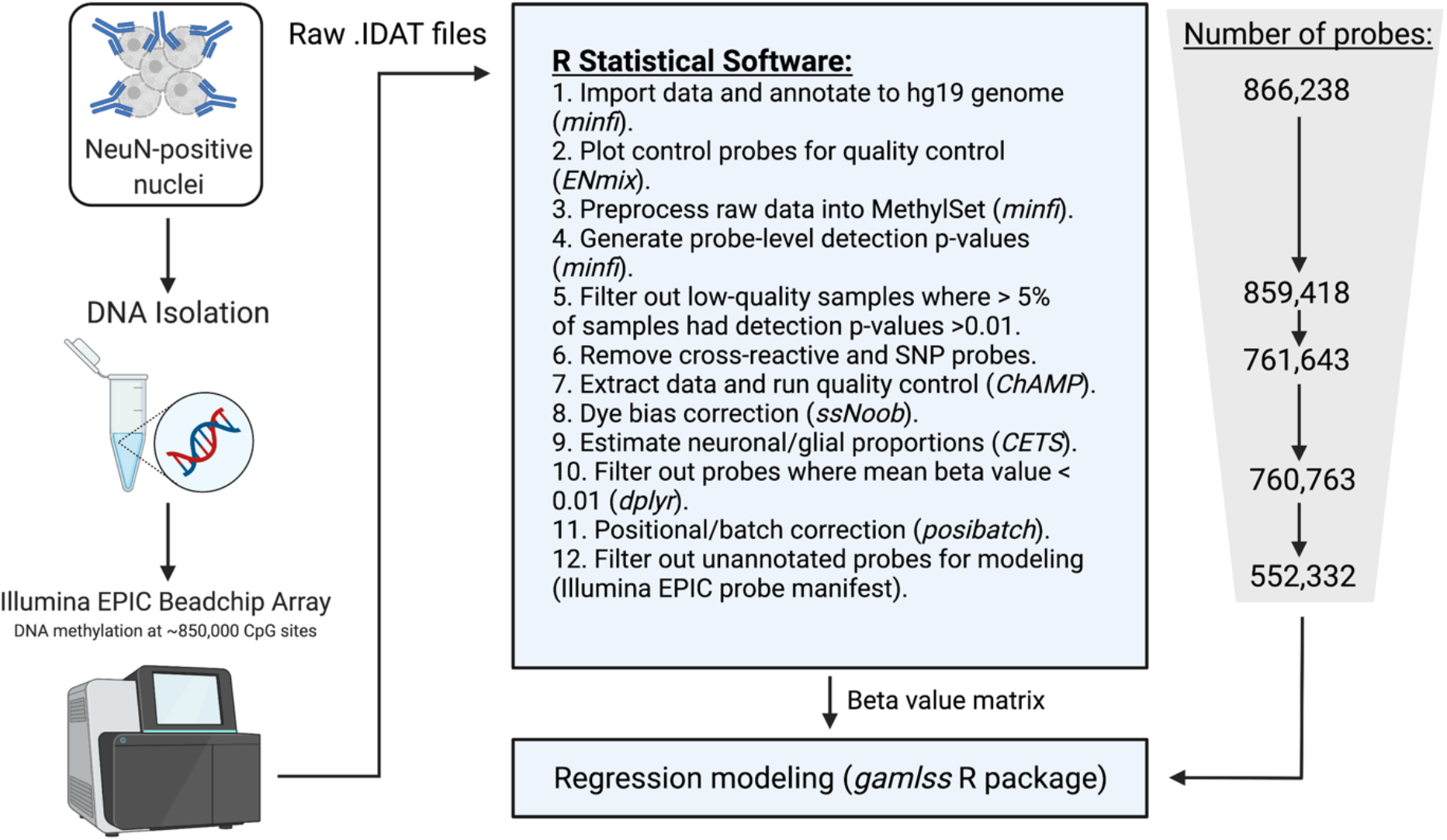
EPIC array data processing pipeline.

### Statistical analysis

#### Differential testing for individual CpG sites

The generalized additive models for location, scale, and shape (*GAMLSS*) R package was used to test for differentially methylated CpGs (DMCs).^92^ This package provides functions to perform multivariate regression modeling while maintaining flexibility regarding term structure and distribution. For DMC testing, we assumed a normal distribution (standard linear regression model) and included both the mu (mean) and sigma (scale parameter) in models to test for differences in mean by disease state while accounting for potential differences in variability. Linear regression was used in lieu of negative-binomial or beta regression modeling due to recent work showing that linear regression does not increase the likelihood of false positives in EPIC array-based epigenome-wide association studies.^48^ All models were stratified by sex, and age and estimated glial cell proportion were included as covariates.

An unadjusted p-value cutoff of < 9×10^-8^ was used to assess significance of DMCs, as recommended in a recent report on human epigenome-wide association studies.^48^ All statistical models were run using R statistical software (version 4.1.0). Annotation of detected differential probes was performed using the Illumina EPIC array manifests. Code for data processing, filtering, and modeling is available as supplementary files (**Supplementary Files 1, 2, and 3**).

#### Differential testing for genomic regions

The *DMRcate* R package was used to test for differentially methylated regions.^93^ In this package, the model matrix was established using the following code: *design = model.matrix(~factor(disease) + glial cell_proportion + age)*. For DMR analysis, Lamba was set to 1000 and the minimum number of CpGs was set to 2. All models were stratified by sex, and both age and estimated glial cell proportion were once again included as covariates in the modeling. The Benjamini-Hochberg False Discovery Rate (FDR) method was used to generate minimum smoothed q-values that account for multiple testing.^94^ For the regional analysis, the significance cutoff was set at a minimum smoothed FDR q-value < 0.05.

#### Gene ontology pathway enrichment

The *clusterProfiler* R package was used to determine whether differentially methylated genes were enriched for specific gene ontology biological process (GOBP) pathways.^95^ Specifically, the enrichGO function was used to run hypergeometric over-representation tests on lists of differentially methylated gene IDs. Gene ID lists were split into hypermethylated DMCs or DMRs and hypomethylated DMCs or DMRs prior to analysis. The list of genes with annotated probes in the EPIC array manifest was used as the universe of total tested genes. Separate, sex-stratified analyses were performed for genes annotated to DMCs and DMRs. GOBP pathway enrichment analysis was not performed on male DMCs due to the low number of significant CpGs (n=3). For enrichment analyses in *clusterProfiler*, redundant pathways were consolidated using the *simplify* function with default parameters, and significance cutoff was set at FDR q-value < 0.05.

#### Epigenetic clock analysis

Epigenetic age was estimated for all parietal cortex samples using a recently published algorithm designed specifically for the human cortex.^82^ Estimates of epigenetic age (years) and known age at death (years) were then compared using standard linear regression with an age*disease interaction term. The interaction term was used to determine whether categorical PD status (control vs. PD) altered trajectories of epigenetic age compared to known chronological age. Epigenetic age regression modeling was performed on all data combined, as well as sex-stratified data. For these regression analyses, significance was set at p-value < 0.05.

## Data availability

This study was preregistered with Open Science Framework: https://osf.io/z4vbw.^96^ All data acquired and analyzed for this study will be deposited in a data repository. Code is also available as Supplementary Material.

## Authors’ contributions

AIB: conceptualization, funding acquisition, methodology, project administration, supervision, writing – review and editing

JK: conceptualization, data curation, formal analysis, investigation, methodology, visualization, writing – original draft, writing – review and editing

NK: conceptualization, investigation, methodology

All authors read and approved the submitted version.

## Acknowledgements

The authors thank the Van Andel Genomics Core, especially Marie Adams, for providing consultation, library preparation, and next-generation sequencing facilities and services (Illumina EPIC array). We also thank the Van Andel Institute Flow Cytometry Core, especially Joshua L. Schipper, for their assistance with flow cytometry. Lastly, we are grateful to the Banner Sun Health Research Institute Brain and Body Donation Program of Sun City, Arizona for the provision of human brain tissue.

