## Supplementary material for "Parkinson’s Disease-Associated, Sex-specific Changes in DNA Methylation at *PARK7* (DJ-1), *ATXN1, SLC17A6, NR4A2*, and *PTPRN2* in Cortical Neurons": biorxiv_Kochmanski_SupplementaryFile1_EPICArray.html

EPIC Array Analysis


Code 

- Show All Code
- Hide All Code

### EPIC Array Analysis

#### 9/2/21

#### Project Summary

This script analyzes BS data from the Illumina EPIC array.

#### Data setup

This assumes you have already downloaded the raw .idat files and prepped a sample sheet, but you can add a step to download the data and create the sample sheet as needed. The sample sheet must include unique sample names in the first column, all relevant data about the samples, and all data from running the arrays, including basename relative to this Rmd document. CETS must be downloaded from Dr. Zachary Kaminsky and the appropriate data for probe masking must also be downloaded from the relevant publication.

- Parent Directory
  - This Rmd document
  - meta\_data.csv (file containing phenotypic data and sample sheet data)
  - idat (folder containing iScan output files)
    - For this project there are 2 subfolders of iScan output
  - CETS\_3.03 (folder contains CETS package, available from Dr. Zachary Kaminsky)
  - Manifest (folder contains manifest and/or files from Zhou et al. 2017 (PMID:27924034))
  - Output (empty folder for output)
    - ChAMP\_Raw (empty folder for ChAMP QC output on raw data)
    - ChAMP\_ssNoob (empty folder for ChAMP QC output on normalized data)
    - Control\_Probes (empty folder for control probe graphs)
  - Illumina manifest file for EPIC is csv format (download available at: https://support.illumina.com/downloads/infinium-methylationepic-v1-0-product-files.html)

#### 1. Load required packages

Load required packages.

```
library(minfi)
library(ChAMP)
library(ENmix)
library(tidyverse)
library(gamlss)
library(parallel)
```

#### 2. Import idat files and set annotation

Set annotation to the correct version of the EPIC array in minfi.

```
# Import meta data
meta <- read.csv("meta_data.csv", header = TRUE)

# Create RGChannelSet objects
rgset <- read.metharray.exp(targets = meta, recursive = TRUE, verbose = TRUE, force = TRUE, extended = TRUE)

# Set annotation
rgset@annotation = c(array='IlluminaHumanMethylationEPIC', annotation='ilm10b4.hg19') #For EPIC array data.

#Check annotation
rgset@annotation

# Save rgsets for easy loading if required
#save(rgset, file = "./Output/rgset.RData")

# If rgsets have already been generated in a previous session, they can be directly imported here.
#load("./Output/rgset.RData")
```

```
##                          array                     annotation 
## "IlluminaHumanMethylationEPIC"                 "ilm10b4.hg19"
```

Data is imported and annotation is set to `rgset@annotation`

#### 3. Check control probes

Check internal control probes with the ENmix function `plotCtrl`.  
Plots are saved in directory ./Output/Control Probes. Review these plots to ensure that the arrays ran correctly.

```
setwd("./Output/Control_Probes/")
dim(rgset)
plotCtrl(rgset)
```

#### 4. Generate MethylSet

Generate MethylSet without normalization using the minfi function `preprocessRaw`.

```
# Set sample names
sampleNames(rgset) <- rgset[[2]]

#Pull out phenotype data
pd <- pData(rgset)

# Generate a MethylSet
mset <- preprocessRaw(rgset)

# Extract the beta values
raw_betas <- getBeta(mset, "Illumina")

# Save mset for easy loading if required
#save(mset, file = "./Output/mset.RData")

# If mset has already been generated in a previous session, they can be directly imported here.
#load("./Output/mset.RData")
```

#### 5. Filter probes and samples

Generate detection p-values and filter out low-quality probes where > 5% of samples had detection p-values >0.01 and samples with greater than 10% of probes failed.

```
# Extract detection p-values
detP <- detectionP(rgset)

# Detect low quality probes based on detection p-value
raw_betas[detP >= 0.01] <- NA

# Calculate the proportion of probes that failed the detection p-value threshold
numfail <- matrix(colMeans(is.na(raw_betas)))

# Rename rows and columns
rownames(numfail) <- colnames(detP)
colnames(numfail) <- "Failed CpG Fraction"
numfail
```

```
##       Failed CpG Fraction
## 00_09        0.0007030400
## 00_27        0.0008404157
## 00_45        0.0025524163
## 01_31        0.0036271787
## 01_37        0.0031584853
## 01_44        0.0020006049
## 01_46        0.0018043540
## 02_15        0.0014245508
## 02_17        0.0015145953
## 02_27        0.0009847178
## 02_33        0.0016554342
## 03_11        0.0007803860
## 03_15        0.0010920786
## 03_25        0.0006383927
## 03_28        0.0008912100
## 03_29        0.0023330771
## 03_39        0.0007192019
## 03_41        0.0011844320
## 03_43        0.0020918039
## 03_47        0.0035440606
## 03_48        0.0011902041
## 03_51        0.0004860096
## 03_63        0.0047481177
## 04_05        0.0017154639
## 04_17        0.0030742129
## 04_38        0.0008750482
## 05_12        0.0016773681
## 05_16        0.0020236933
## 05_17        0.0020363918
## 05_60        0.0013841462
## 06_15        0.0092503446
## 06_21        0.0007769227
## 06_44        0.0017189271
## 06_51        0.0013068002
## 06_57        0.0008623496
## 06_62        0.0014661098
## 07_28        0.0005229510
## 07_37        0.0009835634
## 07_40        0.0069946135
## 07_46        0.0019348031
## 08_40        0.0071169817
## 08_55        0.0016069487
## 08_74        0.0032358313
## 08_85        0.0007665330
## 08_88        0.0008230994
## 08_90        0.0021518336
## 09_12        0.0006245397
## 09_19        0.0005968337
## 09_50        0.0008715849
## 09_52        0.0011994394
## 09_57        0.0007896213
## 10_20        0.0093438524
## 10_22        0.0009824090
## 10_26        0.0014626465
## 10_28        0.0007134298
## 10_39        0.0016081031
## 10_63        0.0020710244
## 10_70        0.0013991536
## 11_54        0.0009166072
## 11_88        0.0005875983
## 11_90        0.0016623607
## 12_21        0.0010886154
## 12_29        0.0009801002
## 12_34        0.0012536970
## 12_42        0.0008115553
## 12_56        0.0015573087
## 13_05        0.0008473422
## 13_11        0.0027625202
## 13_40        0.0009269970
## 13_49        0.0016900667
## 13_52        0.0015261395
## 14_12        0.0007319005
## 15_04        0.0031561765
## 15_46        0.0011902041
## 15_54        0.0011197846
## 15_60        0.0018401409
## 15_69        0.0011728878
## 15_78        0.0009962620
## 16_10        0.0007572976
## 16_23        0.0009927987
## 16_32        0.0007746139
## 17_19        0.0020998848
## 17_37        0.0026620859
## 17_47        0.0025997474
## 17_50        0.0023596286
## 18_05        0.0042782699
## 18_12        0.0030049478
## 18_14        0.0012132924
## 18_17        0.0020537081
## 18_27        0.0010170415
## 18_30        0.0022626576
## 18_33        0.0007515256
## 18_39        0.0020329286
## 18_64        0.0012317631
## 18_65        0.0014857349
## 18_72        0.0017154639
## 18_78        0.0008842835
## 19_02        0.0135055262
## 99_38        0.0013079546
## 99_63        0.0011682702
```

```
#Write .csv of failed probe fractions
write.csv(numfail, "./Output//failed_CpG_fraction.csv")

# Identify samples with greater than 10% of probes failed
RemainSample <- which(numfail < 0.1) 
RemoveSample <- which(numfail > 0.1) 

#Identify and remove failed samples
RemoveSamples <- rownames(numfail)[RemoveSample]                     
meta.r <- meta[RemainSample,]
mset.r <- mset[,RemainSample]
raw_betas.r <- raw_betas[,RemainSample]
detP.r <- detP[,RemainSample]
rgset.r <- rgset[, as.integer(RemainSample)]
pd.r <- pd[RemainSample,]

# Set the probe cutoff to drop probes that failed in > 5% of samples 
ProbeCutoff <- 0.05

# Remove probes that failed in > 5% of samples
mset.f <- mset.r[rowSums(is.na(raw_betas)) <= ProbeCutoff * ncol(detP), ]
raw_betas.f <- raw_betas.r[rowSums(is.na(raw_betas)) <= ProbeCutoff * ncol(detP), ]
detP.f <- detP.r[rowSums(is.na(raw_betas)) <= ProbeCutoff * ncol(detP),]
```

100 of 100 samples remain after filtering samples with a high level (>10%) of failed probes and any corresponding paired samples.  
Of 866238 probes, 859418 remain after removing probes that failed in > 5% of samples.  
6820 probes were removed.

#### 6. Mask probes

Select probe-lists from Zhou for EPIC.

```
# Zhou (for EPIC) - Documentation is found here: http://zwdzwd.github.io/InfiniumAnnotation

# Load files
EPIC.manifest <- readRDS("./Manifest/EPIC.hg19.manifest.rds")
EPIC_manifest_hg19 <- as.data.frame(EPIC.manifest)

# Select probes to mask (see documentation here: http://zwdzwd.github.io/InfiniumAnnotation)
maskname <- rownames(EPIC_manifest_hg19)[which(EPIC_manifest_hg19$MASK_general == TRUE)] 

# Filter mset and raw_betas to remove probes
mset.fm <- mset.f[!featureNames(mset.f) %in% maskname, ]
raw_betas.fm <- raw_betas.f[!rownames(raw_betas.f) %in% maskname, ]
```

After masking, 761643 probes remain. 97775 probes were removed.

#### 7. Adjust beta-values

Fix Beta values that are either 0 or greater than or equal to 1.

```
if (min(raw_betas.fm, na.rm = TRUE) <= 0)
  raw_betas.fm[raw_betas.fm <= 0] <- min(raw_betas.fm[raw_betas.fm > 0])

if (max(raw_betas.fm, na.rm = TRUE) >= 1)
  raw_betas.m.fm[raw_betas.fm >= 1] <- max(raw_betas.fm[raw_betas.fm < 1])
```

Zeros in your dataset have been replaced with smallest positive value.  
Ones in your dataset have been replaced with largest value below 1.

#### 8. Extract raw data and run QC

```
#Get the intensity values and detection p-values using minfi to feed to QC in ChAMP
intensity <- minfi::getMeth(mset.fm) + minfi::getUnmeth(mset.fm)
detP.fm <- detP.r[which(row.names(detP.f) %in% row.names(raw_betas.fm)), ]

#Compile the data into a list object to feed into ChAMP QC
preprocessed.raw.data <- list(mset = mset.f, rgSet = rgset.r, pd = pd.r, intensity = intensity, beta = raw_betas.fm, detP = detP.fm)

#Run QC with ChAMP
champ.QC(beta = preprocessed.raw.data$beta, pheno = pd.r$Disease, resultsDir = "./Output/ChAMP_Raw/")
```

Plots generated by ChampQC can be found in ./Output/ChAMP\_Raw

#### 9. Dye bias correction with ssNoob and run QC

Perform dye bias correction for beta values with ssNoob on probes remaining after filtering and masking.

```
# Perform dye bias correction for beta values with ssNoob on probes remaining after filtering and masking. 
mset.n <- preprocessNoob(rgset.r, dyeMethod = "single")[rownames(raw_betas.fm), ]

# Set sample names
sampleNames(mset.n) = mset.n@colData@rownames

# Extract data from mset
betas.n <- getBeta(mset.n, "Illumina")

#Compile the data into a list object to feed into ChAMP QC
ssNoob.data <- list(mset = mset.n, rgSet = rgset.r, pd = pd.r, intensity = intensity, beta = betas.n, detP = detP.fm)

# Run QC in ChAMP after normalization
#champ.QC(beta = ssNoob.data$beta, pheno = pd$Disease, resultsDir = "./Output/ChAMP_ssNoob/")
```

Plots generated by ChampQC can be found in ./Output/ChAMP\_ssNoob

#### 10. Estimate cell type proportions with CETS

CETS (cell epigenotype specific) estimates neuronal and glial proportions based on methylation data. User should select appropriate controls for CETS.

```
#Load CETS
load("./CETS_3.03/CETS_Image.RData")

# Use glial cell proportion, control brains, Caucasian samples to generate reference 
idx <- list(controlNeuron = pdBrain$celltype == "N" & pdBrain$diag == "Control" & pdBrain$ethnicity == "Caucasian", controlGlia = pdBrain$celltype == "G" & pdBrain$diag == "Control" & pdBrain$ethnicity == "Caucasian")

refProfile <- getReference(brain, idx)

# Estimate neuronal proportion
prop.n <- estProportion(betas.n, profile = refProfile)
round(prop.n, 3)

# Convert to glial proportion (1- neuronal proportion)
prop.n <- as.data.frame(prop.n)
prop.g <- as.data.frame(1 - prop.n) 

# Change rowname to column
prop.n <- tibble::rownames_to_column(prop.n, var = "rowname")
prop.g <- tibble::rownames_to_column(prop.g, var = "rowname")

# Change column name
names(prop.n)[1] <- "Sample"
names(prop.g)[1] <- "Sample"

# Change column name to glial
names(prop.n)[2] <- "neuronal"
names(prop.g)[2] <- "glial"

#Make histogram of results
library(ggplot2)
# Basic histogram
ggplot(prop.n, aes(x=neuronal)) + theme_classic() + geom_histogram(colour="black", fill="white") +
  labs(title="Neuronal proportion histogram plot",x="Neuronal proportion (CETS estimate)", y = "Count")
```

```
# Keep case and glial cell proportion
prop.g$Sample_ID <- prop.g$Sample
keep <- c("Sample_ID", "glial")
prop.g <- prop.g[,keep]

# Add glial cell proportion data to meta data
meta.r$glial <- NA #Create empty variable
meta.r$glial <- prop.g$glial[match(meta.r$Sample_ID, prop.g$Sample_ID)]

write.csv(meta.r, "./meta_data_cets.csv")

# Add glial cell proportion data to pd
pd.r@listData$glial <- NA
pd.r@listData$glial <- prop.g$glial[match(pd.r@listData$Sample_ID, prop.g$Sample_ID)]
```

```
## 00_09 00_27 00_45 01_31 01_37 01_44 01_46 02_15 02_17 02_27 02_33 03_11 03_15 
## 0.919 0.921 0.903 0.923 0.806 0.415 0.679 0.556 0.755 0.991 0.631 0.533 0.770 
## 03_25 03_28 03_29 03_39 03_41 03_43 03_47 03_48 03_51 03_63 04_05 04_17 04_38 
## 0.751 0.639 0.805 0.855 0.919 0.834 0.919 0.928 0.030 0.739 0.476 0.595 0.469 
## 05_12 05_16 05_17 05_60 06_15 06_21 06_44 06_51 06_57 06_62 07_28 07_37 07_40 
## 0.959 0.974 0.921 0.804 0.746 0.562 0.812 0.883 0.640 0.736 0.649 1.000 0.942 
## 07_46 08_40 08_55 08_74 08_85 08_88 08_90 09_12 09_19 09_50 09_52 09_57 10_20 
## 1.000 0.994 0.961 0.552 0.641 0.850 0.906 0.752 0.604 0.984 0.937 0.706 0.719 
## 10_22 10_26 10_28 10_39 10_63 10_70 11_54 11_88 11_90 12_21 12_29 12_34 12_42 
## 0.772 0.891 1.000 0.977 0.843 1.000 0.970 0.904 0.891 0.940 0.906 0.757 0.967 
## 12_56 13_05 13_11 13_40 13_49 13_52 14_12 15_04 15_46 15_54 15_60 15_69 15_78 
## 0.792 1.000 1.000 0.924 1.000 0.919 0.801 1.000 0.784 0.950 1.000 0.766 0.955 
## 16_10 16_23 16_32 17_19 17_37 17_47 17_50 18_05 18_12 18_14 18_17 18_27 18_30 
## 0.998 0.808 0.936 0.799 0.845 0.817 0.933 0.803 0.837 0.946 0.771 0.915 0.851 
## 18_33 18_39 18_64 18_65 18_72 18_78 19_02 99_38 99_63 
## 0.881 0.898 1.000 0.741 0.946 0.959 0.902 0.833 0.869
```

#### 11. Run ChAMP SVD

Run SVD to determine potential batch effects.

```
champ.SVD(beta = betas.n, pd = pd.r, resultsDir = "./Output/CHAMP_SVD_BS/")
```

```
##       Sample_ID Subject.ID   subject     Title     Index Sample_Well
##  [1,] 0.4810969  0.4810969 0.4810969 0.4810969 0.4810969   0.9200780
##  [2,] 0.4810969  0.4810969 0.4810969 0.4810969 0.4810969   0.3129493
##  [3,] 0.4810969  0.4810969 0.4810969 0.4810969 0.4810969   0.4151122
##  [4,] 0.4810969  0.4810969 0.4810969 0.4810969 0.4810969   0.2724906
##  [5,] 0.4810969  0.4810969 0.4810969 0.4810969 0.4810969   0.1644510
##  [6,] 0.4810969  0.4810969 0.4810969 0.4810969 0.4810969   0.4694275
##  [7,] 0.4810969  0.4810969 0.4810969 0.4810969 0.4810969   0.2272818
##  [8,] 0.4810969  0.4810969 0.4810969 0.4810969 0.4810969   0.2021768
##  [9,] 0.4810969  0.4810969 0.4810969 0.4810969 0.4810969   0.6451075
##       Sample_Plate    Disease Disease_coded   Sentrix_ID       Array
##  [1,] 1.082713e-04 0.88489230    0.88489230 3.949411e-05 0.188590301
##  [2,] 1.922057e-05 0.27605532    0.27605532 1.406201e-02 0.003432419
##  [3,] 1.096119e-01 0.66406287    0.66406287 7.640359e-01 0.805055944
##  [4,] 9.809860e-02 0.86858848    0.86858848 4.169200e-04 0.042916900
##  [5,] 6.134475e-05 0.44005057    0.44005057 3.307523e-01 0.014174828
##  [6,] 1.545707e-01 0.06772048    0.06772048 3.790194e-01 0.040901476
##  [7,] 7.952556e-03 1.00000000    1.00000000 1.558416e-03 0.843067497
##  [8,] 5.513833e-01 0.56719526    0.56719526 8.384522e-02 0.033554639
##  [9,] 3.105549e-01 0.13110811    0.13110811 8.671358e-01 0.328343354
##              Batch         Date      Race       Gender        Age       PMI
##  [1,] 1.817727e-03 1.817727e-03 0.0929236 4.011491e-02 0.41897986 0.1207537
##  [2,] 8.898132e-05 8.898132e-05 0.6649898 1.627918e-01 0.70363582 0.6760960
##  [3,] 1.109925e-01 1.109925e-01 0.7420755 1.177477e-15 0.23772518 0.1539006
##  [4,] 5.006444e-02 5.006444e-02 0.9861802 9.572976e-01 0.75590756 0.9345697
##  [5,] 2.137759e-05 2.137759e-05 0.9585573 8.667605e-01 0.44998970 0.4612235
##  [6,] 5.510886e-02 5.510886e-02 0.8761179 6.298725e-01 0.24142371 0.4733291
##  [7,] 1.807425e-01 1.807425e-01 0.2751644 8.723806e-01 0.16687851 0.7107418
##  [8,] 4.945848e-01 4.945848e-01 0.8488898 9.800647e-01 0.03804851 0.1830143
##  [9,] 1.676272e-01 1.676272e-01 0.2602124 6.919324e-01 0.80672980 0.2768106
##           Location        glial
##  [1,] 1.817727e-03 3.886254e-71
##  [2,] 8.898132e-05 1.420439e-01
##  [3,] 1.109925e-01 5.454897e-01
##  [4,] 5.006444e-02 8.512545e-01
##  [5,] 2.137759e-05 9.443681e-01
##  [6,] 5.510886e-02 5.834839e-01
##  [7,] 1.807425e-01 9.547884e-01
##  [8,] 4.945848e-01 8.662386e-01
##  [9,] 1.676272e-01 6.439378e-01
```

SVD Summary files can be found in ./Output/CHAMP\_SVD\_BS

#### 12. Remove outliers based on neuron/glial cell proportion estimates

After reviewing CETS output, select a cutoff as necessary.

```
#Remove samples with high (>0.90) proportion of glial cells. Select this cutoff as necessary based on CETS results
Keep <- prop.g[prop.g$glial <0.9,1]
Remove <- prop.g[prop.g$glial >0.9,1]
mc <- as.data.frame(betas.n[,Keep])

#Remove samples from meta data (where SampleID is in Keep)
meta.r.o <- meta.r[meta.r$Sample_ID %in% Keep,]

#Write .csv of the limited meta data.
write.csv(meta.r.o, "./meta_data_cets_outlier_removed.csv")
```

1 samples have been removed due to low neuronal content. Meta data has been saved with outliers removed.

Samples removed: 03\_51

```
#Save estimated beta values:
save(mc, file = "./EPIC_betas.RData")

#After saving, can load back in easily:
#load("./EPIC_betas.RData")
```

Estimated beta values have been saved as an RData file.

Your beta values (EPIC\_betas.RData) and meta data (meta\_data\_cets\_outlier\_removed.csv) are ready for differential methylation testing!
