## Supplementary material for "Parkinson’s Disease-Associated, Sex-specific Changes in DNA Methylation at *PARK7* (DJ-1), *ATXN1, SLC17A6, NR4A2*, and *PTPRN2* in Cortical Neurons": biorxiv_Kochmanski_SupplementaryFile2_EPICArray.Rmd

````
---
title: "EPIC Array Analysis"
authors: Joe Kochmanski, Alison Bernstein
date: "9/2/21"
output:
  html_document:
    toc: yes
    toc_float: yes
    df_print: paged
    code_folding: hide
    fig_height: 2.5
    fig_width: 3.5
    highlight: tango
  html_notebook:
    code_folding: hide
    fig_height: 2.5
    fig_width: 3.5
    highlight: tango
    toc: yes
    toc_float: yes
editor_options:
  chunk_output_type: console
---

```{r setup, include=FALSE}
library(knitr)
knitr::opts_chunk$set(echo = TRUE, cache.lazy = FALSE)
```

## Project Summary
This script analyzes BS data from the Illumina EPIC array.

## Data setup
This assumes you have already downloaded the raw .idat files and prepped a sample sheet, but you can add a step to download the data and create the sample sheet as needed. The sample sheet must include unique sample names in the first column, all relevant data about the samples, and all data from running the arrays, including basename relative to this Rmd document.
CETS must be downloaded from Dr. Zachary Kaminsky and the appropriate data for probe masking must also be downloaded from the relevant publication.

* Parent Directory
  + This Rmd document
  + meta_data.csv (file containing phenotypic data and sample sheet data)
  + idat (folder containing iScan output files)
      - For this project there are 2 subfolders of iScan output
  + CETS_3.03 (folder contains CETS package, available from Dr. Zachary Kaminsky)
  + Manifest (folder contains manifest and/or files from Zhou et al. 2017 (PMID:27924034))
  + Output (empty folder for output)
    - ChAMP_Raw (empty folder for ChAMP QC output on raw data)
    - ChAMP_ssNoob (empty folder for ChAMP QC output on normalized data)
    - Control_Probes (empty folder for control probe graphs)
  + Illumina manifest file for EPIC is csv format (download available at: https://support.illumina.com/downloads/infinium-methylationepic-v1-0-product-files.html)


## 1. Load required packages
Load required packages.
```{r Load packages, echo=TRUE, warning=FALSE, message=FALSE, cache=FALSE, results='hide' }
library(minfi)
library(ChAMP)
library(ENmix)
library(tidyverse)
library(gamlss)
library(parallel)
```

## 2. Import idat files and set annotation
Set annotation to the correct version of the EPIC array in minfi. 
```{r Create RGChannelSet, echo=TRUE, warning=FALSE, message=FALSE, cache=TRUE, results='hold', cache=FALSE}
# Import meta data
meta <- read.csv("meta_data.csv", header = TRUE)

# Create RGChannelSet objects
rgset <- read.metharray.exp(targets = meta, recursive = TRUE, verbose = TRUE, force = TRUE, extended = TRUE)

# Set annotation
rgset@annotation = c(array='IlluminaHumanMethylationEPIC', annotation='ilm10b4.hg19') #For EPIC array data.

#Check annotation
rgset@annotation

# Save rgsets for easy loading if required
#save(rgset, file = "./Output/rgset.RData")

# If rgsets have already been generated in a previous session, they can be directly imported here.
#load("./Output/rgset.RData")

```
Data is imported and annotation is set to `rgset@annotation`

## 3. Check control probes
Check internal control probes with the ENmix function `plotCtrl`.  
Plots are saved in directory ./Output/Control Probes. Review these plots to ensure that the arrays ran correctly. 
```{r Internal control probes, echo=TRUE, warning=FALSE, message=FALSE, cache=TRUE, results='hide'}
setwd("./Output/Control_Probes/")
dim(rgset)
plotCtrl(rgset) 
```

## 4. Generate MethylSet
Generate MethylSet without normalization using the minfi function `preprocessRaw`.
```{r Generate MethylSet, echo=TRUE, warning=FALSE, message=FALSE, cache=TRUE, results='hold'}
# Set sample names
sampleNames(rgset) <- rgset[[2]]

#Pull out phenotype data
pd <- pData(rgset)

# Generate a MethylSet
mset <- preprocessRaw(rgset)

# Extract the beta values
raw_betas <- getBeta(mset, "Illumina")

# Save mset for easy loading if required
#save(mset, file = "./Output/mset.RData")

# If mset has already been generated in a previous session, they can be directly imported here.
#load("./Output/mset.RData")
```

## 5. Filter probes and samples
Generate detection p-values and filter out low-quality probes where > 5% of samples had detection p-values >0.01 and samples with greater than 10% of probes failed.
```{r Filter data by detection p-value, echo=TRUE, warning=FALSE, message=FALSE, cache=TRUE}

# Extract detection p-values
detP <- detectionP(rgset)

# Detect low quality probes based on detection p-value
raw_betas[detP >= 0.01] <- NA

# Calculate the proportion of probes that failed the detection p-value threshold
numfail <- matrix(colMeans(is.na(raw_betas)))

# Rename rows and columns
rownames(numfail) <- colnames(detP)
colnames(numfail) <- "Failed CpG Fraction"
numfail

#Write .csv of failed probe fractions
write.csv(numfail, "./Output//failed_CpG_fraction.csv")

# Identify samples with greater than 10% of probes failed
RemainSample <- which(numfail < 0.1) 
RemoveSample <- which(numfail > 0.1) 

#Identify and remove failed samples
RemoveSamples <- rownames(numfail)[RemoveSample]                     
meta.r <- meta[RemainSample,]
mset.r <- mset[,RemainSample]
raw_betas.r <- raw_betas[,RemainSample]
detP.r <- detP[,RemainSample]
rgset.r <- rgset[, as.integer(RemainSample)]
pd.r <- pd[RemainSample,]

# Set the probe cutoff to drop probes that failed in > 5% of samples 
ProbeCutoff <- 0.05

# Remove probes that failed in > 5% of samples
mset.f <- mset.r[rowSums(is.na(raw_betas)) <= ProbeCutoff * ncol(detP), ]
raw_betas.f <- raw_betas.r[rowSums(is.na(raw_betas)) <= ProbeCutoff * ncol(detP), ]
detP.f <- detP.r[rowSums(is.na(raw_betas)) <= ProbeCutoff * ncol(detP),]
```
`r toString(nrow(meta.r))` of `r toString(nrow(meta))` samples remain after filtering samples with a high level (>10%) of failed probes and any corresponding paired samples.  
Of `r toString(length(mset@NAMES))` probes, `r toString(length(mset.f@NAMES))` remain after removing probes that failed in  > `r toString(ProbeCutoff*100)`% of samples.  
`r toString(length(mset@NAMES)-length(mset.f@NAMES))` probes were removed.

## 6. Mask probes
Select probe-lists from Zhou for EPIC.
```{r Mask probes, echo=TRUE, warning=FALSE, message=FALSE, cache=TRUE}
# Zhou (for EPIC) - Documentation is found here: http://zwdzwd.github.io/InfiniumAnnotation

# Load files
EPIC.manifest <- readRDS("./Manifest/EPIC.hg19.manifest.rds")
EPIC_manifest_hg19 <- as.data.frame(EPIC.manifest)

# Select probes to mask (see documentation here: http://zwdzwd.github.io/InfiniumAnnotation)
maskname <- rownames(EPIC_manifest_hg19)[which(EPIC_manifest_hg19$MASK_general == TRUE)] 

# Filter mset and raw_betas to remove probes
mset.fm <- mset.f[!featureNames(mset.f) %in% maskname, ]
raw_betas.fm <- raw_betas.f[!rownames(raw_betas.f) %in% maskname, ]
```
After masking, `r toString(length(mset.fm@NAMES))` probes remain. 
`r toString(length(mset.f@NAMES)-length(mset.fm@NAMES))` probes were removed.

## 7. Adjust beta-values
Fix Beta values that are either 0 or greater than or equal to 1.
```{r Fix Beta values, echo=TRUE, warning=FALSE, message=FALSE, cache=TRUE}
if (min(raw_betas.fm, na.rm = TRUE) <= 0)
  raw_betas.fm[raw_betas.fm <= 0] <- min(raw_betas.fm[raw_betas.fm > 0])

if (max(raw_betas.fm, na.rm = TRUE) >= 1)
  raw_betas.m.fm[raw_betas.fm >= 1] <- max(raw_betas.fm[raw_betas.fm < 1])
```
Zeros in your dataset have been replaced with smallest positive value.  
Ones in your dataset have been replaced with largest value below 1.  

## 8. Extract raw data and run QC
```{r Extract raw data and QC, echo=TRUE, warning=FALSE, message=FALSE, cache=TRUE}
#Get the intensity values and detection p-values using minfi to feed to QC in ChAMP
intensity <- minfi::getMeth(mset.fm) + minfi::getUnmeth(mset.fm)
detP.fm <- detP.r[which(row.names(detP.f) %in% row.names(raw_betas.fm)), ]

#Compile the data into a list object to feed into ChAMP QC
preprocessed.raw.data <- list(mset = mset.f, rgSet = rgset.r, pd = pd.r, intensity = intensity, beta = raw_betas.fm, detP = detP.fm)

#Run QC with ChAMP
champ.QC(beta = preprocessed.raw.data$beta, pheno = pd.r$Disease, resultsDir = "./Output/ChAMP_Raw/")
```
Plots generated by ChampQC can be found in ./Output/ChAMP_Raw

## 9. Dye bias correction with ssNoob and run QC
Perform dye bias correction for beta values with ssNoob on probes remaining after filtering and masking. 
```{r Dye bias correction, echo=TRUE, warning=FALSE, message=FALSE, cache=TRUE, results='hide'}

# Perform dye bias correction for beta values with ssNoob on probes remaining after filtering and masking. 
mset.n <- preprocessNoob(rgset.r, dyeMethod = "single")[rownames(raw_betas.fm), ]

# Set sample names
sampleNames(mset.n) = mset.n@colData@rownames

# Extract data from mset
betas.n <- getBeta(mset.n, "Illumina")

#Compile the data into a list object to feed into ChAMP QC
ssNoob.data <- list(mset = mset.n, rgSet = rgset.r, pd = pd.r, intensity = intensity, beta = betas.n, detP = detP.fm)

# Run QC in ChAMP after normalization
#champ.QC(beta = ssNoob.data$beta, pheno = pd$Disease, resultsDir = "./Output/ChAMP_ssNoob/")
```
Plots generated by ChampQC can be found in ./Output/ChAMP_ssNoob

## 10. Estimate cell type proportions with CETS
CETS (cell epigenotype specific) estimates neuronal and glial proportions based on methylation data. User should select appropriate controls for CETS.
```{r CETS, echo=TRUE, warning=FALSE, message=FALSE, cache=TRUE, results='hold'}

#Load CETS
load("./CETS_3.03/CETS_Image.RData")

# Use glial cell proportion, control brains, Caucasian samples to generate reference 
idx <- list(controlNeuron = pdBrain$celltype == "N" & pdBrain$diag == "Control" & pdBrain$ethnicity == "Caucasian", controlGlia = pdBrain$celltype == "G" & pdBrain$diag == "Control" & pdBrain$ethnicity == "Caucasian")

refProfile <- getReference(brain, idx)

# Estimate neuronal proportion
prop.n <- estProportion(betas.n, profile = refProfile)
round(prop.n, 3)

# Convert to glial proportion (1- neuronal proportion)
prop.n <- as.data.frame(prop.n)
prop.g <- as.data.frame(1 - prop.n) 

# Change rowname to column
prop.n <- tibble::rownames_to_column(prop.n, var = "rowname")
prop.g <- tibble::rownames_to_column(prop.g, var = "rowname")

# Change column name
names(prop.n)[1] <- "Sample"
names(prop.g)[1] <- "Sample"

# Change column name to glial
names(prop.n)[2] <- "neuronal"
names(prop.g)[2] <- "glial"

#Make histogram of results
library(ggplot2)
# Basic histogram
ggplot(prop.n, aes(x=neuronal)) + theme_classic() + geom_histogram(colour="black", fill="white") +
  labs(title="Neuronal proportion histogram plot",x="Neuronal proportion (CETS estimate)", y = "Count")

# Keep case and glial cell proportion
prop.g$Sample_ID <- prop.g$Sample
keep <- c("Sample_ID", "glial")
prop.g <- prop.g[,keep]

# Add glial cell proportion data to meta data
meta.r$glial <- NA #Create empty variable
meta.r$glial <- prop.g$glial[match(meta.r$Sample_ID, prop.g$Sample_ID)]

write.csv(meta.r, "./meta_data_cets.csv")

# Add glial cell proportion data to pd
pd.r@listData$glial <- NA
pd.r@listData$glial <- prop.g$glial[match(pd.r@listData$Sample_ID, prop.g$Sample_ID)]
```

## 11. Run ChAMP SVD
Run SVD to determine potential batch effects.
```{r Run ChAMP on raw BS beta values with glial cell content, echo=TRUE, warning=FALSE, message=FALSE, cache=TRUE, fig.width=7, fig.height=5}
champ.SVD(beta = betas.n, pd = pd.r, resultsDir = "./Output/CHAMP_SVD_BS/")
```
SVD Summary files can be found in ./Output/CHAMP_SVD_BS

## 12. Remove outliers based on neuron/glial cell proportion estimates
After reviewing CETS output, select a cutoff as necessary.
```{r Remove outliers, echo=TRUE, warning=FALSE, message=FALSE, cache=TRUE}

#Remove samples with high (>0.90) proportion of glial cells. Select this cutoff as necessary based on CETS results
Keep <- prop.g[prop.g$glial <0.9,1]
Remove <- prop.g[prop.g$glial >0.9,1]
mc <- as.data.frame(betas.n[,Keep])

#Remove samples from meta data (where SampleID is in Keep)
meta.r.o <- meta.r[meta.r$Sample_ID %in% Keep,]

#Write .csv of the limited meta data.
write.csv(meta.r.o, "./meta_data_cets_outlier_removed.csv")
```
`r toString(ncol(betas.n)-ncol(mc))` samples have been removed due to low neuronal content. Meta data has been saved with outliers removed.

Samples removed: `r toString(Remove) ` 

```{r Save final beta values as Rdata, echo=TRUE, warning=FALSE, message=FALSE, cache=TRUE}
#Save estimated beta values:
save(mc, file = "./EPIC_betas.RData")

#After saving, can load back in easily:
#load("./EPIC_betas.RData")

```
Estimated beta values have been saved as an RData file.

Your beta values (EPIC_betas.RData) and meta data (meta_data_cets_outlier_removed.csv) are ready for differential methylation testing!
````
