## Supplementary material for "Parkinson’s Disease-Associated, Sex-specific Changes in DNA Methylation at *PARK7* (DJ-1), *ATXN1, SLC17A6, NR4A2*, and *PTPRN2* in Cortical Neurons": biorxiv_Kochmanski_SupplementaryFile4_080921.pdf

1.fcs
Ungated
31716

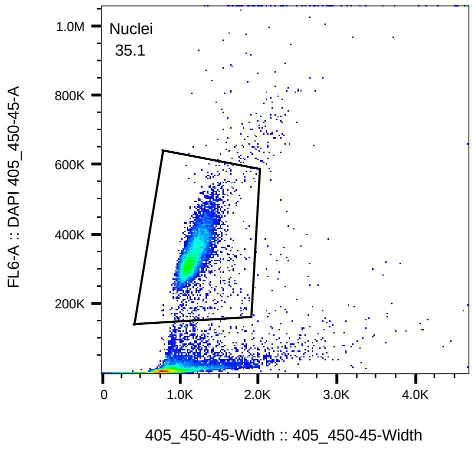


1.fcs
Nuclei
11131

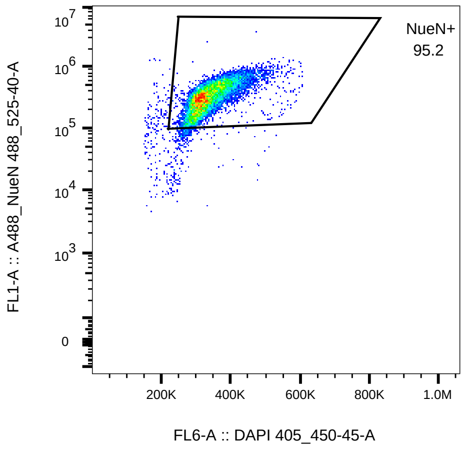


2.fcs
Ungated
28546

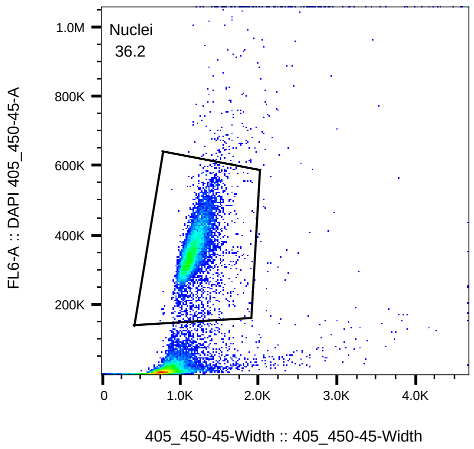


2.fcs
Nuclei
10330

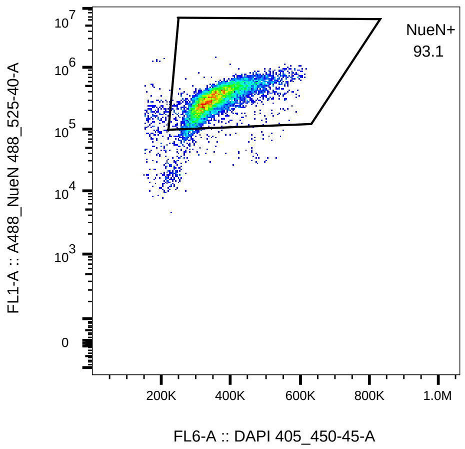


3.fcs
Ungated
30666

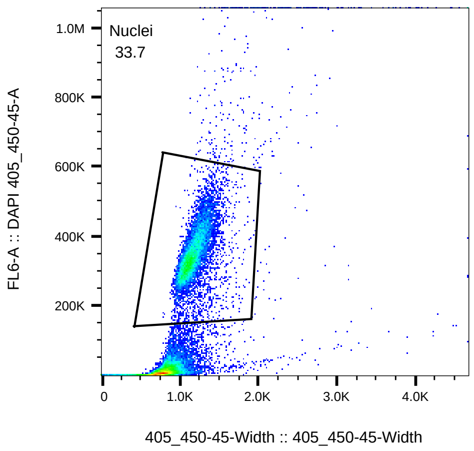


3.fcs
Nuclei
10344

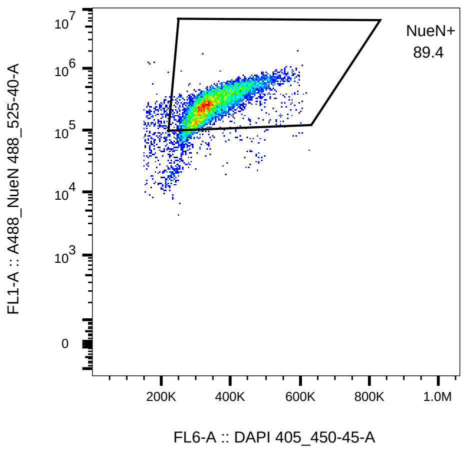

4.fcs
Ungated
22807

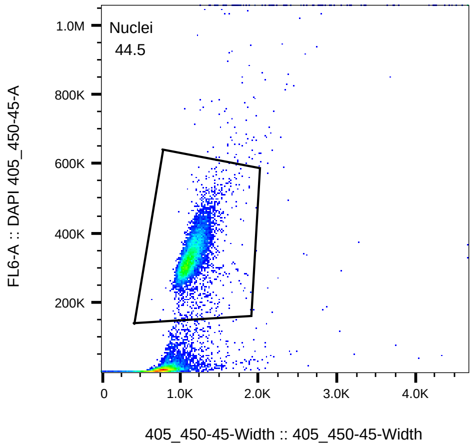


4.fcs
Nuclei
10146

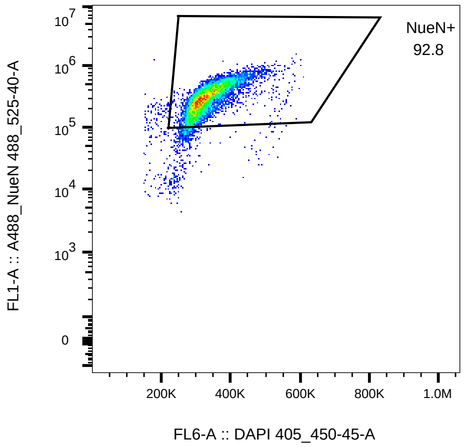


5.fcs
Ungated
12403

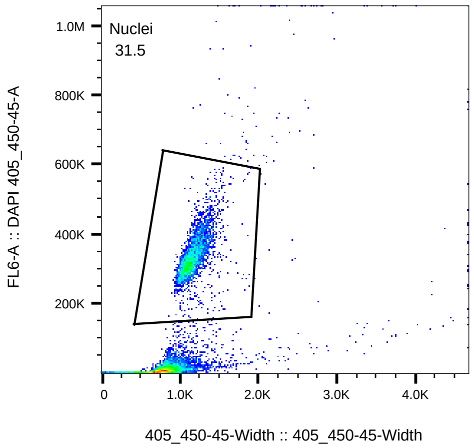


5.fcs
Nuclei
3903

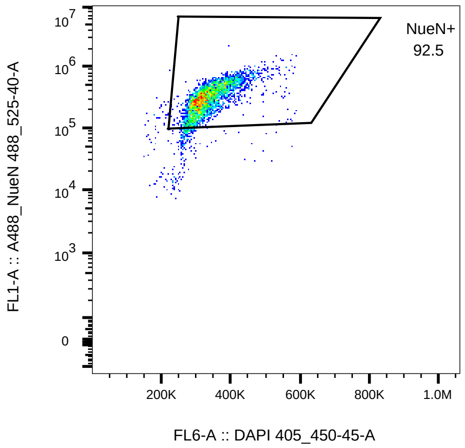


6.fcs
Ungated
22811

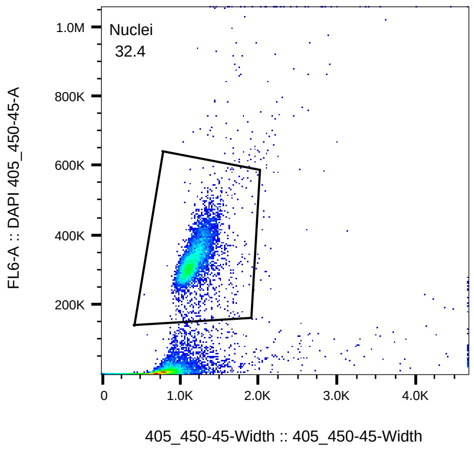


6.fcs
Nuclei
7394

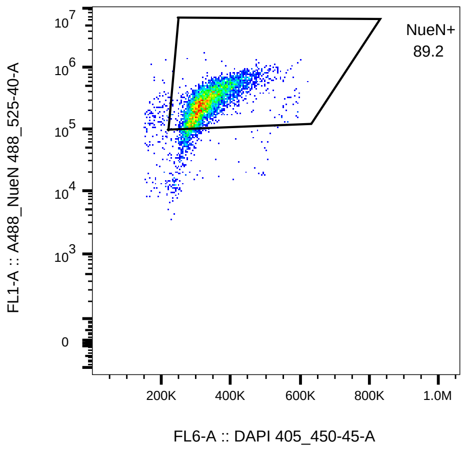

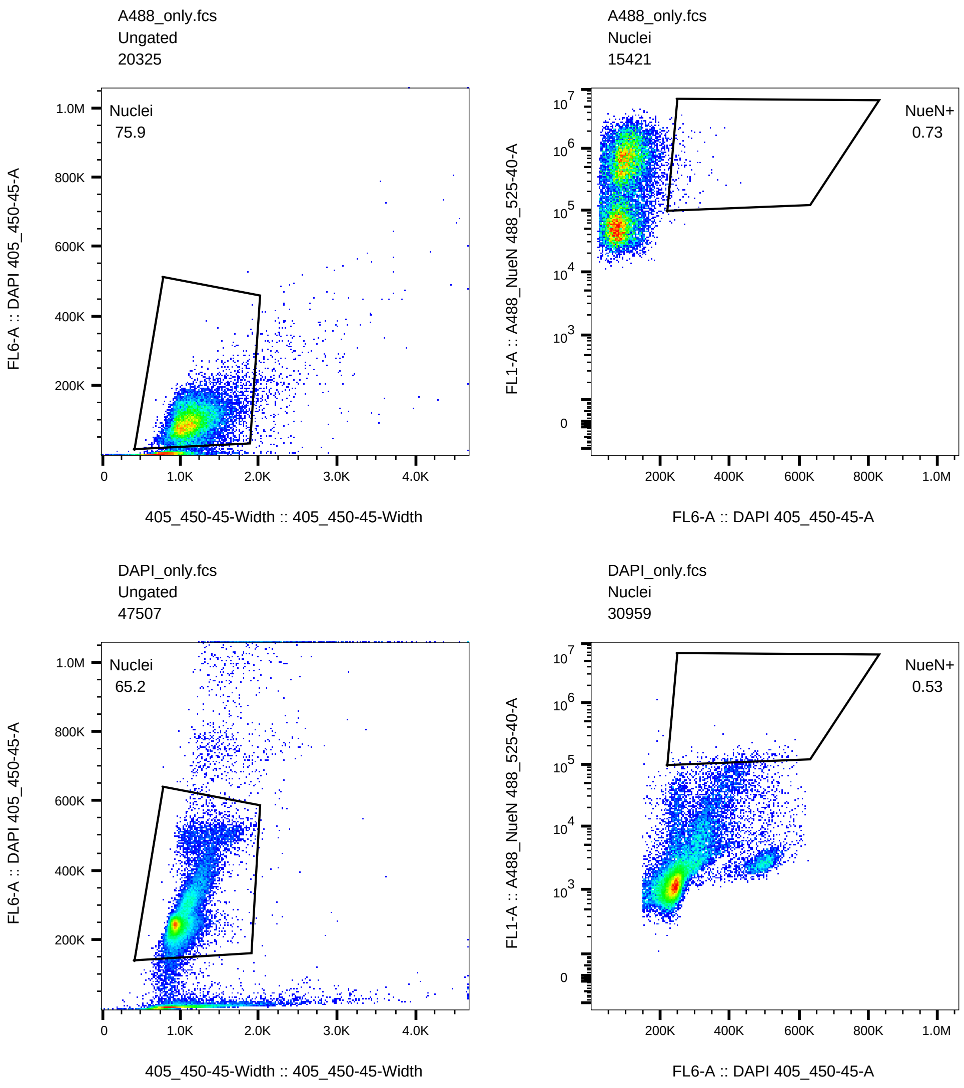
A488_only.fcs
Ungated
20325
Nuclei
75.9
FL6-A :: DAPI 405_450-45-A
405_450-45-Width :: 405_450-45-Width
A488_only.fcs
Nuclei
15421
NueN+
0.73
FL1-A :: A488_NueN 488_525-40-A
FL6-A :: DAPI 405_450-45-A
DAPI_only.fcs
Ungated
47507
Nuclei
65.2
FL6-A :: DAPI 405_450-45-A
405_450-45-Width :: 405_450-45-Width
DAPI_only.fcs
Nuclei
30959
NueN+
0.53
FL1-A :: A488_NueN 488_525-40-A
FL6-A :: DAPI 405_450-45-A
